# *miR-200* deficiency promotes lung cancer metastasis by activating cancer-associated fibroblasts

**DOI:** 10.1101/2020.09.02.276550

**Authors:** Bin Xue, Chen-Hua Chuang, Haydn M. Prosser, Cesar Seigi Fuziwara, Claudia Chan, Neil Sahasrabudhe, Maximilian Kühn, Yalei Wu, Jingqi Chen, Anne Biton, Caifu Chen, John Erby Wilkinson, Michael T. McManus, Allan Bradley, Monte M Winslow, Bo Su, Lin He

## Abstract

Lung adenocarcinoma, the most prevalent lung cancer subtype, is characterized by its high propensity to metastasize. Despite the importance of metastasis in lung cancer mortality, its underlying cellular and molecular mechanisms remain largely elusive. Here, we identified *miR-200* miRNAs as potent suppressors for lung adenocarcinoma metastasis. *miR-200* expression is specifically repressed in mouse metastatic lung adenocarcinomas, and *miR-200* decrease strongly correlates with poor patient survival. Consistently, deletion of *mir-200c/141* in the *Kras^LSL-G12D/+^; Trp53^flox/flox^* lung adenocarcinoma mouse model significantly promoted metastasis, generating a desmoplastic tumor stroma highly reminiscent of metastatic human lung cancer. *miR-200* deficiency in lung cancer cells promotes the proliferation and activation of adjacent cancer-associated fibroblasts (CAFs), which in turn elevates the metastatic potential of cancer cells. *miR-200* regulates the functional interaction between cancer cells and CAFs, at least in part, by targeting Notch ligand Jagged1 and Jagged2 in cancer cells and inducing Notch activation in adjacent CAFs. Hence, the interaction between cancer cells and CAFs constitutes an essential mechanism to promote metastatic potential.

## Introduction

Lung adenocarcinoma, the most prevalent lung cancer subtype, is characterized by its high propensity to metastasize. Metastatic lung adenocarcinoma cells escape primary tumors, disseminate via blood and lymphatic circulation, and ultimately colonize distant organs, particularly the liver, bone and brain^1,2^. This metastasis results in rapid disease progression and organ failure, ultimately accounting for the majority of patient mortality^3^. Despite its immense clinical significance, the cellular and molecular basis for lung adenocarcinoma metastasis remains largely unknown.

Both cancer cell intrinsic mechanisms and tumor-stroma interactions regulate metastatic progression^4^. Previous studies have mostly focused on intrinsic mechanisms that promote metastasis, involving transcription factors and signaling molecules that increase chromosome instability and accessibility, promote developmental plasticity, alter the cancer secretome, or enhance epithelial-mesenchymal transition (EMT)^5–8^. However, emerging evidence has implicated interactions between cancer cells and the tumor microenvironment, as lung adenocarcinoma metastases are frequently characterized by desmoplasia and substantial remodeling of the tumor microenvironment^9,10^. One major obstacle to study lung adenocarcinoma metastasis is the limited number of faithful animal models that recapitulates the entire metastatic processes *in vivo*. Widely used transplantation models largely depend on cancer cell lines, often bypassing important *in vivo* processes to achieve metastatic growth^11^, while popular genetically engineered mouse models (GEMMs) of lung adenocarcinoma often require a relatively long latency to develop metastasis^12,13^.

One of the most widely used models for metastatic lung adenocarcinoma is the *Kras^LSL-G12D/+^; Trp53^flox/flox^* (*KP*) model, in which inducible *Kras^G12D^* expression and *p53* deletion initiate the growth of lung adenocarcinomas that highly resemble human pathology^14^. While this model can eventually manifest metastatic tumors, its usefulness is hampered by a long latency period and incomplete penetrance^12,13^. Here, we identified *miR-200* miRNAs as key suppressors for metastasis in *KP* lung adenocarcinomas. Engineered *miR-200* deficiency promoted the rapid development of metastasis to lymph nodes and distant organs, faithfully recapitulating the desmoplastic tumor microenvironment characteristic of metastatic human disease. *miR-200* deficiency in neoplastic cells increases the expression of Notch ligand Jag1 and Jag2. Enhanced Notch signaling in neighboring stromal fibroblasts induced their proliferation and their remodeling into cancer-associated fibroblasts (CAFs), which in turn promoted the development of metastases. Our findings uncover key miRNAs that normally suppress metastasis by restricting CAF activation, and highlights the importance of tumor microenvironment remodeling during lung adenocarcinoma metastasis.

## Results

### Downregulation of *miR-200* miRNAs in a mouse model for metastatic lung cancer

To compare the miRNA expression profiles between primary and metastatic lung adenocarcinomas, we employed the *Kras*^LSL-*G12D/+*^*;Trp53^flox/flox^;R26^LSL-tdTomato/+^* (*KPT*) lung adenocarcinoma mouse model, in which inducible *Kras* activation and *p53* loss in lung epithelial cells yield malignant lung adenocarcinomas with a long latency for metastasis^12,13^. Spontaneous metastasis emerged in lymph nodes, peritoneum, liver, adrenal gland, and soft tissue 5 to 9 months after tumor initiation induced by lenti-Cre administration^13,15^. *KPT* lung cancer cells were marked with the tdTomato Cre-reporter, which enabled purification of cancer cells from primary tumors and metastases (Fig. 1A). Among 641 miRNAs analyzed, 20 highly-expressed miRNAs were differentially expressed between non-metastatic primary tumors and metastases (Fig. 1B, Sup Fig. S1A, Supplementary Table S1). Strikingly, nearly all *miR-200* miRNAs were repressed in metastases (Sup Fig. S1A). In particular, *miR-141* and *miR-200c* were among the most down-regulated miRNAs in metastatic *KPT* lung tumors (Fig. 1B, Sup Fig. S1A).

**Figure 1.**
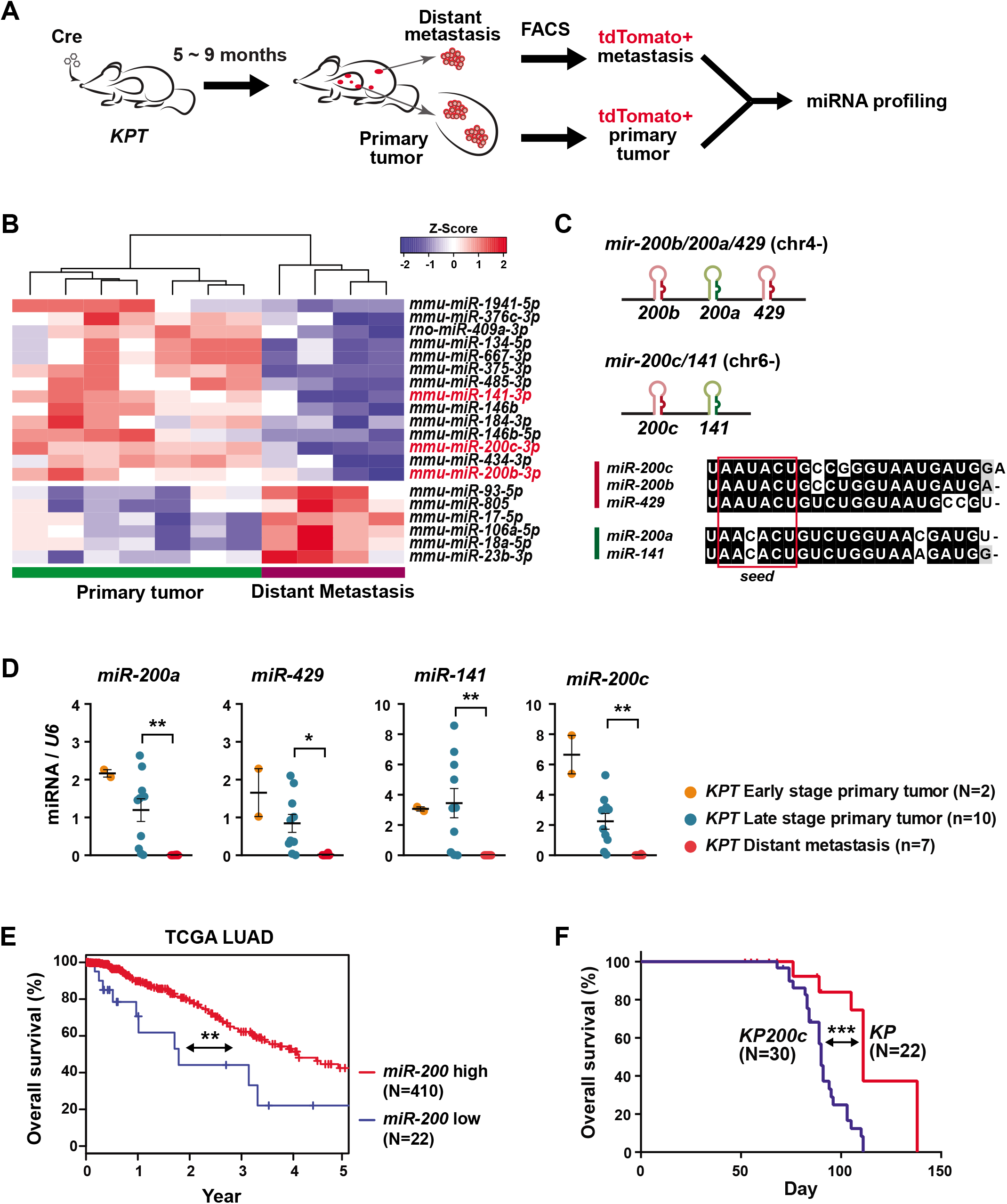
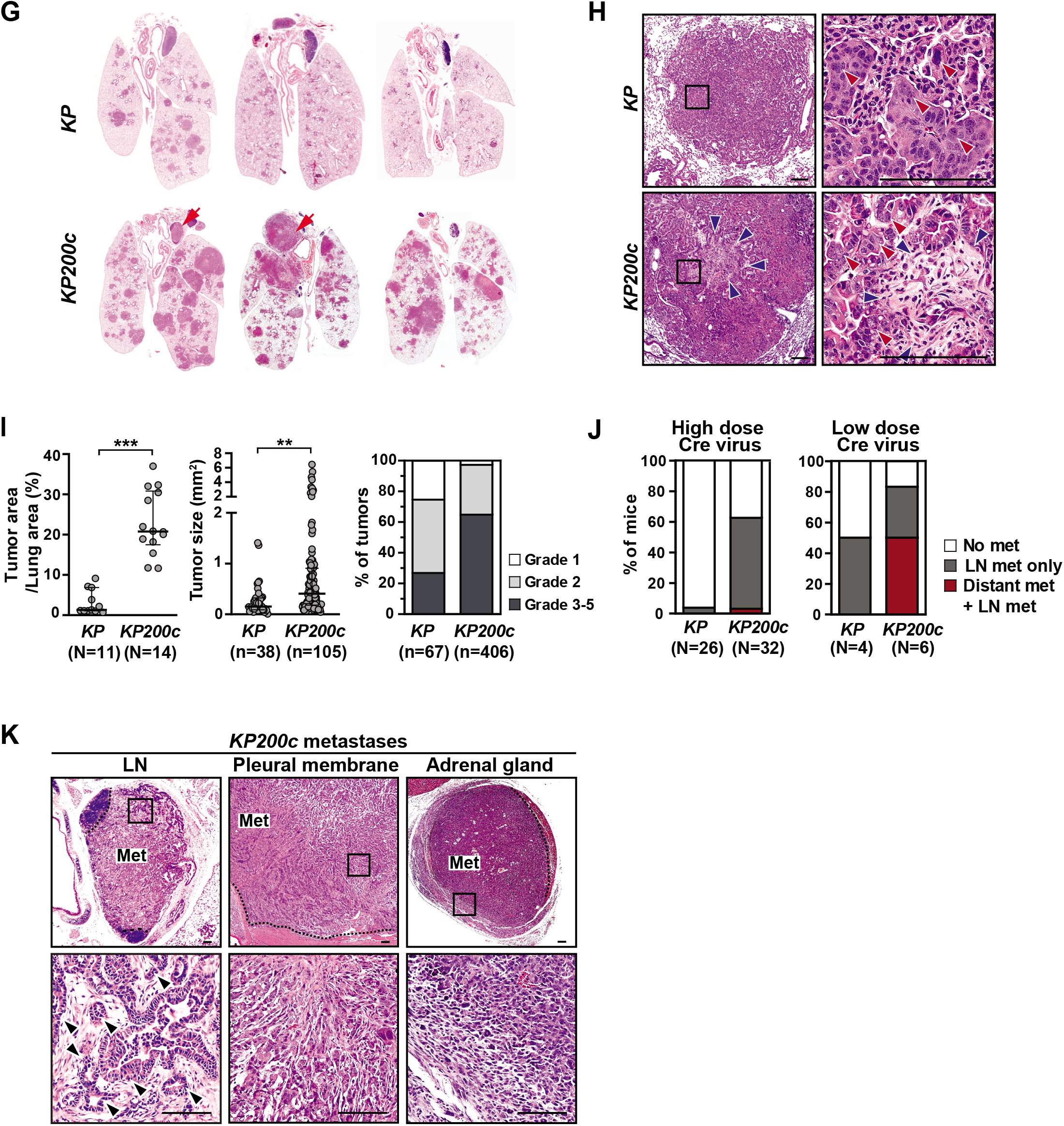
*miR-200* deficiency induces rapid metastasis in *KP* lung cancer model. **A.** Cartoon illustrating isolation of tdTomato+ cancer cells from primary and metastatic *KPT* lung adenocarcinomas for miRNA profiling. **B.** *miR-200* miRNAs are strongly downregulated in distant metastases in the *KPT* mouse lung adenocarcinoma model. A Heatmap is shown for the most differentially expressed miRNAs between seven *KPT* primary lung tumors and four distant metastases (*P*<0.05 & Expression level ≥−20). Red text, *miR-200* miRNAs. **C.** The five *miR-200* miRNAs reside in two genomic loci and segregate into two sub-families based on seed sequences. Red box, seed sequences. **D.** *miR-200* miRNAs are strongly downregulated in metastatic *KPT* tumors. Real time PCR detected *miR-200* expression in *KP* early-stage primary tumor (N=2) and late-stage primary tumor (n=10), but not in *KP* metastases (n=7). Error bars, s.e.m.; late stage primary tumors vs. metastases: *miR-200a*, ** *P*=0.0052, t=3.268, df=15; *miR-429:* * *P*=0.0113, t=2.887, df=15; *miR-141:* ** *P*=0.01, t=2.047, df=15; *miR-200c:* ** *P*=0.0068, t=3.395, df=10. All statistical analyses were performed using unpaired, two tailed, student’s t-test. **E.** Decreased total *MIR-200* expression human lung adenocarcinomas is associated with poor patient survival in the TCGA data (LUAD). A Kaplan-Meier plot compares patient survival between two cohort of patients, with either high (N=410) or low (N=22) expression of all five *MIR-200* miRNAs. (** *P*= 0.00188, log rank test). **F.** *miR-200c/141* deficiency in the *KP200c* model significantly reduces overall survival. A Kaplan-Meier plot compares survival of *KP* and *KP200c* mice after Adeno-Cre administration (5×10^6^ Ad-Cre PFU/mouse), demonstrating a significant acceleration of tumor progression in *KP200c* mice. *** *P*= 0.0003, log rank test. **G.** *miR-200c/141* deficiency in *KP200c* mice induces a significant increase in tumor progression. Representative H&E staining of lung sections are shown for three pairs of *KP* and *KP200c* mice collected at 90 days after tumor initiation. Red arrows, lymph node metastases. **H.** *KP200c* primary tumors are characterized by desmoplastic stroma (blue arrows) and pleomorphic nuclei in cancer cells (red arrows). Representative images are shown for a pair of late-stage *KP* and *KP200c* primary tumors collected at terminal stage. Scale bars, 100μm **I.** *miR-200* deficiency leads to a significant increase in tumor burden, tumor size, and tumor grade in *KP200c* mice. Tumor burden is quantified as the percentage of tumor area versus total lung area (left, error bars, s.e.m. *** *P*<0.0001, N, animal number, unpaired two-tailed Student’s t-test, t=7.7222, df=23); tumor size was measured by tumor area (middle, error bars, s.e.m., \*\**P*=0.006, n, tumor number, unpaired two-tailed Student’s t-test, t=2.792, df=141); tumor grade was determined by histopathological scores (right, n, tumor number. *** *P*<0.0001, χ^2^=79.43, df=2). All *KP* and *KP200c* tumors were collected at 12 weeks after tumor initiation. **J.** *KP200c* mice exhibit a greater metastasis frequency compared to *KP* mice. 26 *KP* and 32 *KP200c* mice were compared upon tumor initiation with high-dose Cre virus (5×10^6^ Ad-Cre PFU/mouse, *** *P*<0.0001, χ^2^=149.3, df=2), with *KP200c* mice developing more LN metastases. 4 *KP* and 6 *KP200c* mice were compared upon tumor initiation with low-dose Cre virus (1×10^5^ Lenti-Cre PFU/mouse, *** *P*<0.0001, χ^2^=2428, df=2), with *KP200c* mice developing more distant metastases. **K.** *KP200c* mice develop metastases in multiple sites. Representative images of H&E staining of a lymph node metastasis, a pleural metastasis, and a distant metastasis in adrenal gland from *KP200c* mice are shown. Dotted line shows the boundary of metastasis and normal tissue. Scale bar, 100μm.

The *miR-200* family contains five evolutionarily conserved miRNAs located in two distinct genomic loci, *mir-200b/200a/429* on chromosome 4 and *mir-200c/141* on chromosome 6. They are classified into two subfamilies that differ by one nucleotide in their seed sequences (Fig. 1C). *miR-200* miRNAs were expressed in both early and late stage primary lung adenocarcinomas, exhibiting only a slight downregulation during tumor progression (Fig. 1D). However, *miR-200* expression was strongly repressed specifically in metastatic lung adenocarcinomas in *KPT* mice (Fig. 1D). Consistently, in the 432 lung adenocarcinomas analyzed by The Cancer Genome Atlas (TCGA) project, a collective, low *miR-200* expression was correlated with poor overall patient survival (\*\*\**P*=0.0018, Fig. 1E). Hence, downregulation of *miR-200* is associated with tumor metastasis in lung adenocarcinomas.

### *miR-200* deficiency promotes the development of lung cancer metastases

We next investigated the functional role of *miR-200* miRNAs in lung cancer metastasis in the *Kras*^LSL-*G12D/+*^*;Trp53^flox/flox^* (*KP*) mouse model^13,14^. We engineered individual deletion of *mir-200c/141* or *mir-200b/200a/429* in mice^16^ (Sup Fig. S1B). *mir-200c/141^-/-^* mice were phenotypically normal, with no obvious developmental defects. However, deficiency of *mir-200b/200a/429* impaired fertility^17^, while *mir-200c/141^-/-^; mir-200b/200a/429^-/-^* mice are neonatal lethal (data not shown). While further deletion of *mir-200c/141* in *KP mice* (*KP;mir-200c/141^-/-^*, designated as the *KP200c*) exhibit only partial *miR-200* deficiency, they nonetheless provide essential insights into the role of *miR-200* in lung cancer metastases.

Upon tumor initiation with Adeno-Cre virus (5×10^6^ Ad-Cre PFU/mouse), *KP200c* mice quickly developed highly malignant lung adenocarcinomas. Advanced lesions with severe pleomorphic nuclei were present as early as one month after tumor initiation (Sup Fig. S1C). At the same time point, *KP* mice only had adenomas with mild nuclei atypia (Sup Fig. S1C). In line with these observations, *KP200c* mice exhibited a significant decrease in overall survival compared to *KP* mice (*KP200c* median survival 90 days, *KP* median survival 111 days, *** *P*=0.0003, log rank test, Fig. 1F, Supplementary Table S2).

At 12 weeks after tumor initiation, *KP200c* mice developed significantly larger and more aggressive lung adenocarcinomas, with a tumor burden 5 times as high as that of *KP* mice (Fig. 1G, 1I). Nearly all *KP200c* lung tumors were grade 2 or above, with more than 65% between grade 3 to grade 5, while 25% of *KP* tumors were grade 1 and only 27% were grade 3 or above (Fig. 1I). High grade *KP200c* tumors were characterized by their large size, solid histological pattern, large and pleomorphic nuclei, and high degree of desmoplastic stroma surrounding the cancer cells (Fig. 1H).

*miR-200c/141* deficiency not only accelerated tumor growth and progression, but also significantly increased the frequency of tumor metastasis. More than 60% of *KP200c* mice (20 out of 32 mice) developed lymph node (LN) metastases within 120 days, mostly occurring in mediastinal and thoracic lymph nodes (Fig. 1G, Fig 1J, Supplementary Table S2). Within the same time frame, only 1 in 26 *KP* mice developed LN metastasis (Fig 1J). When a lower dose of Cre-virus was used to initiate tumors in *KP200c* and *KP* mice (1×10^5^ Lenti-Cre PFU/mouse), 50% of *KP200c* mice developed full-blown distant metastases to liver, pleura, pericardium, and adrenal gland by 150 days after tumor initiation (Fig. 1J, Fig. 1K, Sup Fig. S1E, Supplementary Table S2). Under the same experimental conditions, *KP* mice exhibited much lower primary tumor burden with no distant metastasis (Fig. 1J, Supplementary Table S2). Interestingly, all *KP200c* metastases exhibited a complete silencing of *mir-200b/200b/429* (Sup Fig. S1D), suggesting a strong selective pressure for a complete *miR-200* loss in metastatic *Kras*-driven, *p53*-null lung adenocarcinomas. Hence, *KP200c* mice provide a powerful new experimental system to probe the underlying mechanisms of adenocarcinoma metastasis.

### *miR-200* deficiency induces the expansion of CAFs in metastatic *KP200c* tumors

Metastases in *KP200c* mice were accompanied by an increase in MAPK signaling and a loss of cell differentiation (Fig. 2A). Compared to *KP* primary tumors, phosphorylated Erk1/2 level was significantly increased in *KP200c* primary tumors (\**P*=0.0263), and further elevated in *KP200c* LN and distant metastases (\*\*\**P*<0.0001) (Fig. 2A), indicating a synergic effect of *Kras^G12D^* induction and *mir-200c/141* deficiency on MAPK activation during metastasis progression. In addition, 64% of lymph node and distant metastases from *KP200c* mice were negative for both surfactant protein C (SPC), a marker for type 2 alveolar epithelial cells (AT2), and club cell antigen 10 (CC10), a marker for bronchiolar club cells (Fig. 2B). *KP200c* primary tumors, in comparison, were more heterogeneous in the expression of SPC and CC10, with only 28% of tumors being SPC^low^CC10^low^, and the remaining exhibiting varying degrees of SPC and CC10 expression (Fig. 2B, Sup Fig. S2A). Nearly all *KP200c* metastases lost expression of Nkx2.1 (Fig. 2C), a lung lineage transcription factor that controls lung tumor differentiation and limits metastatic potential^6^. *KP200c* primary tumors, in comparison, mostly expressed Nkx2.1 with downregulated expression in some high-grade regions (Fig. 2C). Our findings suggest that *KP200c* metastases could be derived from poorly differentiated cancer cells with highly elevated MAPK signaling.

**Figure 2.**
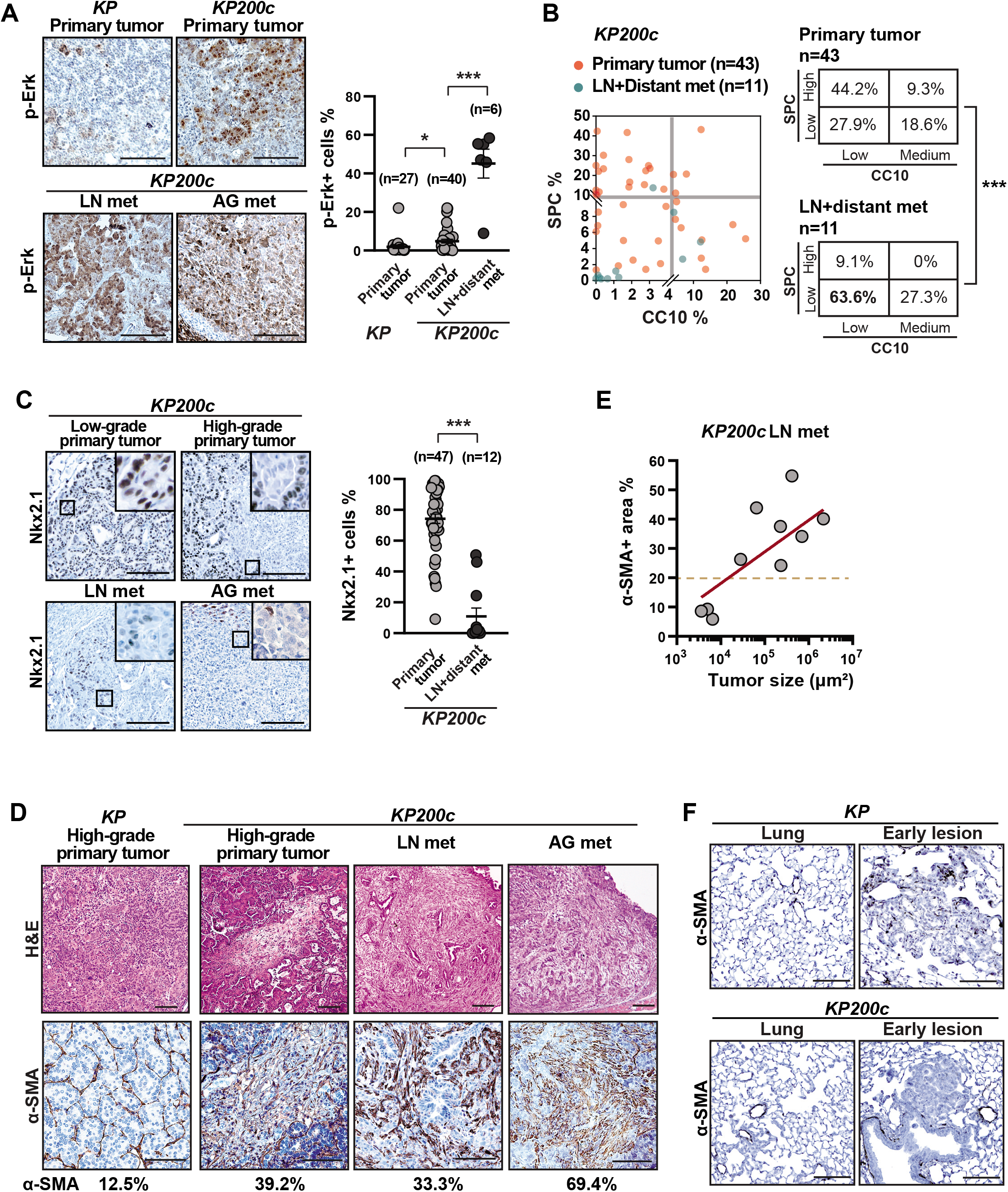
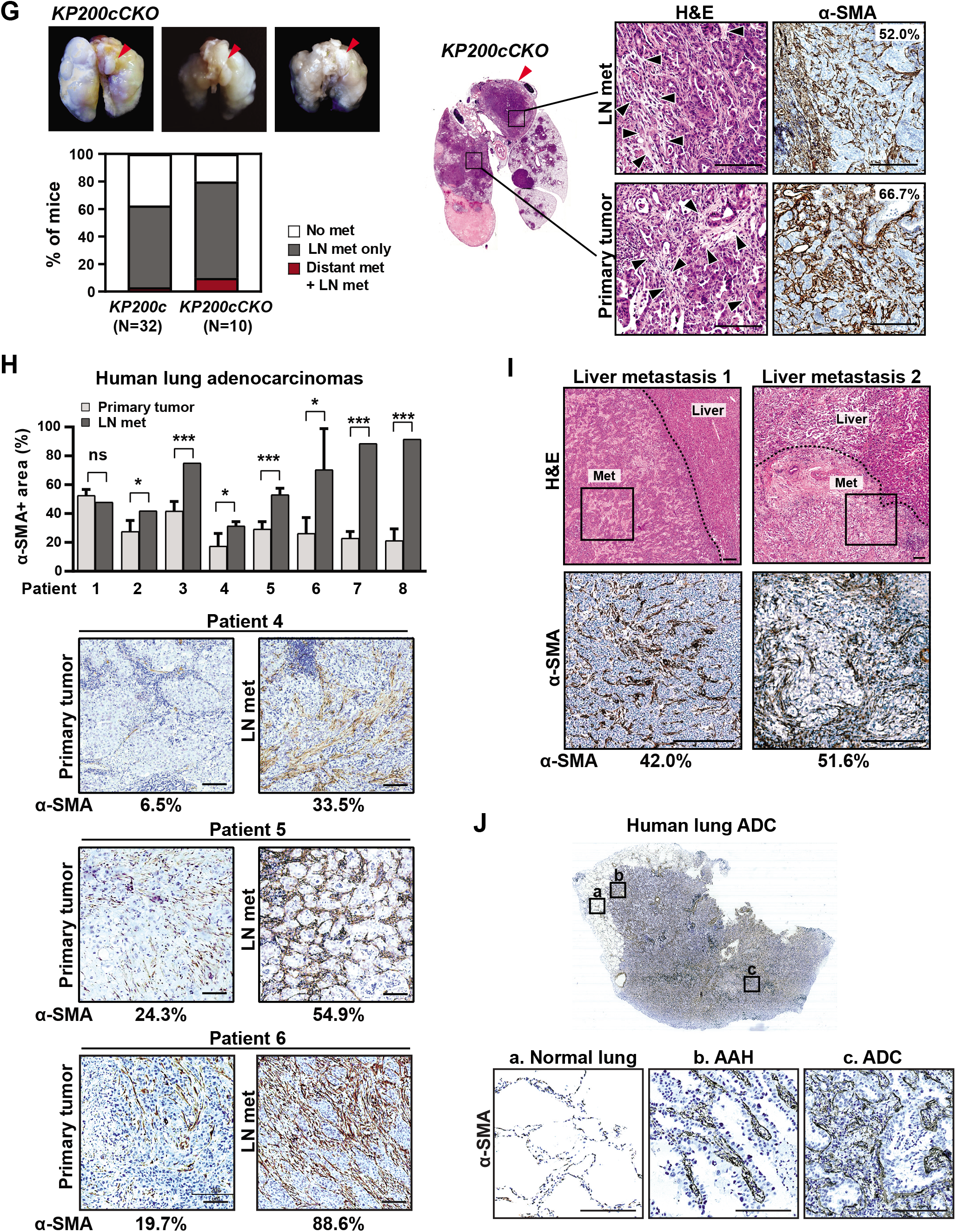
CAFs are enriched in metastatic *KP200c* lung adenocarcinomas. **A.** Primary and metastatic *KP200c* tumors exhibit a strong increase in MAPK signaling. Immunohistochemistry staining of phosphorylated Erk1/2 (p-Erk) (left) and quantitation of p-Erk-positive cells (right) indicate a strong elevation of MAPK signaling in *KP200c* tumors, particularly the metastatic *KP200c* tumors. 27 *KP* primary tumors, 40 *KP200c* primary tumors and 6 *KP200c* metastases were analyzed. Scale bar, 100μm. Error bar, s.e.m., *KP* vs. *KP200c* primary tumor, \**P*=0.0263, unpaired two-tailed Student’s t-test, t=2.273, df=65; *KP200c* primary vs *KP200c* lymph node metastases, \*\*\**P*<0.0001, unpaired two-tailed Student’s t-test, t=11.06, df=44. **B.** Metastatic *KP200c* tumors are characterized by a loss of cell lineage markers. Quantitation is shown for immunohistochemistry staining of CC10 (a marker for Club cells) and SPC (a marker for alveolar type 2 cells) in *KP200c* primary and metastatic tumors (left). Tumor stratification shows a preferential loss of CC10 and SPC expression in *KP200c* metastases compared to *KP200c* primary tumors. *KP200c* primary tumors vs. metastases *** *P*<0.0001, χ^2^=61.67, df=3. **C.** Metastatic *KP200c* tumors exhibit a downregulation of Nkx2. Representative images (left) and quantitation (right) of Nkx2.1 immunohistochemistry staining are shown for *KP200c* primary tumors, lymph node, and distant metastases. Scale bar, 100μm. Error bar, s.e.m., \*\*\**P*<0.0001, unpaired two-tailed Student’s t-test, t=9.474, df=57. 47 *KP200c* primary tumors and 12 *KP200c* metastases (both lymph nodeand distant metastases) were analyzed**. D, E.** α-SMA-expressing CAFs are enriched in *KP200c* primary tumors and metastases. **D.** Representative images of H&E and α-SMA staining are shown high grade regions of *KP* and *KP200c* primary tumors, as well as *KP200c* metastases. Scale bars, 100μm. **E.** Abundance of α-SMA+ CAFs is correlated with the size of *KP200c* metastases. The percentage of α-SMA staining in total tumor area is plotted in a linear regression against the size of metastatic tumors for 10 *KP200c* lymph node metastases, *R*^2^=0.3599. **F.** α-SMA positive fibroblasts are absent from *KP* and *KP200c* normal lung and early primary tumors. Representative images of α-SMA staining are shown for *KP* and *KP200c* normal lung and early primary tumors. Scale bars, 100μm. **G.** Deficiency of *miR-200c/141* specifically in cancer cells promotes metastasis progression and desmoplastic stroma. Left, *KP200cCKO* lungs collected 90 days after tumor initiation (5×10^6^ Adeno-Cre PFU/mouse) exhibit evident lymph node metastases, with metastasis frequency comparable to that of *KP* mice. Right, H&E and α-SMA staining of high grade regions of *KP200cCKO* primary tumors and lymph node metastases indicate strong CAF enrichment in the tumor microenvironment. Scale bars, 100μm. **H, I.** Human metastatic lung adenocarcinomas exhibit an increase of α-SMA+ CAFs compared to paired primary tumors. **H.** Quantitation of α-SMA+ area was shown for 8 pairs of human primary and metastatic lung adenocarcinomas; representative immunohistochemistry staining images are shown for three pairs (Scale bars, 100μm). Pair 2, * *P*=0.0148, t=4.101, df=4.000; pair 3 *** *P*=0.0004, t=11.22, df=4.000; pair 4 * *P*=0.0467, t=2.883, df=3.869; pair 5, *** *P*=0.0010, t=6.955, df=4.951; pair 6 * *P*=0.0250, t=3.530, df=3.920; pair 7 *** *P*=0.0001, t=27.80, df=3.000; pair 8 *** *P*<0.0001, t=19.42, df=4.000. All statistical analyses were performed using unpaired two-tailed Student’s t-test with Welch’s correction. **I.** α-SMA+ CAFs are enriched in liver metastases from relapsed human lung adenocarcinoma patients. H&E staining are shown for two independent liver metastases samples and α-SMA staining are shown for the boxed metastasis region. Dotted line indicates the tumor/liver boundary. Scale bars, 100μm. **J**. Increased cancer-CAF interaction is associated with tumor progression in human lung adenocarcinomas. A stage IA lung adenocarcinoma was stained for α-SMA, regions representing a progressive histopathological pattern including normal lung, atypical adenomatous hyperplasia (AAH), and adenocarcinoma (ADC) are marked and zoom in. Scale bar, 100μm.

A prominent cellular hallmark of metastatic *KP200c* tumors was the abundance of collagen-rich, stromal desmoplasia within the metastatic tumor microenvironment. In comparison, *KP* tumors only exhibited widespread desmoplasia in liver metastases, but not in the lymph node metastases nor in primary tumors (Sup Fig. S2B, S2C, S2D). A major source of extracellular matrix in the tumor microenvironment came from CAFs, a heterogeneous population of fibroblasts that promote cancer growth, survival, and invasiveness by secreting cytokines and growth factors (interleukins, HGF, CXCLs, TGF-β etc.)^18–24^, as well as collagen to increase tissue stiffness^25^. In *KP200c* late-stage primary tumors, lymph node metastases, and distant metastases, we observed a marked increase of α-smooth muscle actin-expressing CAFs (α-SMA+ CAFs) (Fig. 2D), which grew into multiple layers and intertwined with clusters of *KP200c* cancer cells (Fig. 2D). This multilayered, α-SMA+, stromal structure was largely absent in *KP* primary tumors, including high-grade tumors (Fig. 2D). Unlike α-SMA+ CAFs, FSP-1-expressing CAFs did not show a significant increase in *KP200c* tumors (Sup. Fig S2E). Importantly, the percentage of α-SMA+ CAFs in *KP200c* lymph node metastasis was proportional to the tumor size (Fig. 2E), with 70% of *KP200c* lymph node metastases containing at least 20% of α-SMA+ CAFs (Fig. 2E). α-SMA staining was absent from the epithelial compartment in both *KP* and *KP200c* normal lung and early lesions (Fig. 2F). Our findings suggest that *miR-200* deficiency in lung adenocarcinomas lead to the recruitment and expansion of α-SMA+ CAFs in the tumor microenvironment, which could be a critical step for the development of metastatic lung adenocarcinomas.

In *KP200c* mice, both neoplastic cells and stromal cells were deficient for *mir-200c/141*, making it difficult to determine whether *mir-200c/141* deficiency in cancer cells and/or stromal cells drove tumor growth and metastasis. To address the role of mir-200c/141 specifically in neoplastic cells, we generated *KP* mice with a *mir-200c/141* conditional allele (*Kras^G12D/+^; p53^flox/flox^; mir-200c/141^flox/-^*, designated as *KP200cCKO* mice)^26^. This model allows the generation of *mir-200c/141*-deficient *KP* cancer cells in an otherwise *mir-200c/141*-expressing tumor microenvironment (Sup Fig. S2F). Similar to *KP200c* mice, *KP200cCKO* mice exhibited an increase in metastasis frequency, as 80% of *KP200cCKO* mice developed lymph node or distant metastases by 120 days after tumor initiation (Fig. 2G, Supplementary Table S2, Sup Fig. S2G). Metastases in *KP200cCKO* mice resembled those of *KP200c* mice; both were characterized by strong desmoplasia with extensive infiltration of α-SMA+ CAFs (Fig. 2G). Hence, *miR-200* deficiency in lung cancer cells promotes local and distant metastases, while inducing the expansion of α-SMA+ CAFs in the tumor microenvironment.

### Enrichment of SMA+ CAFs occurs in human metastatic lung adenocarcinomas

Consistent with α-SMA+ CAF enrichment in metastatic *KP200c* mouse lung tumors, α-SMA+ CAFs were also enriched in metastatic human lung adenocarcinomas (Fig. 2H). We compared SMA staining in paired primary lung adenocarcinomas and lymph node metastases from 8 patients, and we observed a significant increase of α-SMA+ CAFs in lymph node metastases in 7 out of 8 patients (Fig. 2H). Several patients exhibited at least a two-fold increase in the number of α-SMA+ CAFs in lymph node metastases compared to their paired primary tumors (Fig. 2H). Two patients exhibited a similar enrichment of α-SMA+ CAFs in liver metastases relative to their paired primary tumors (Fig. 2I).

Interestingly, we captured a continuum of CAF activation patterns in a stage IA human lung adenocarcinoma (Fig. 2J). In normal lung, the thin alveoli septa consists of a monolayer of α-SMA+ pericytes embedded in the basement membrane of type 1 alveolar cells (AT1) (Fig. 2J, area a). When the lesion progressed to atypical adenomatous hyperplasia (AAH), the transformed epithelial cells grew into the alveolar space with the α-SMA+ CAFs starting to expand within the alveolar septa (Fig. 2J, area b). In high-grade adenocarcinomas where the alveolar lumen is filled with cancer cells (Fig. 2J, area c), the α-SMA+ CAFs formed multilayered structures intermingled with cancer cells. Hence, the expansion of the α-SMA+ CAFs in the tumor microenvironment, as well as the extent of tumor-CAF interaction, both correlate with histological progression in human adenocarcinomas.

### Fibroblast co-culture promotes the metastatic features in *KP200c* tumor organoids

We next established a 3D tumor organoid system by culturing one *KP* and three *KP200c* lung cancer cell lines (Sup Fig. S3A) on top of a matrix containing 50% Matrigel+0.5mg/ml Collagen I to mimic a collagen-rich lung tumor microenvironment (Fig. 3A). When cultured alone, *KP* and *KP200c* cancer cells both formed well-polarized spherical tumor organoids with hollow lumina (Fig. 3B), with *KP200c* organoids slightly larger in size (Fig. 3B), but with neither exhibiting invasive features (Fig. 3B). Interestingly, when co-cultured with primary lung fibroblasts, the miR-200 deficient *KP200c* organoids, but not the miR-200 expressing-*KP* organoids, clustered to form large multi-acinar structures with cellular protrusions, indicative of invasive behavior (Fig. 3C, 3D). This clustering behavior substantially increased the size of *KP200c* tumor organoids in culture (Fig. 3B, Sup Fig. S3B). The extent of organoid clustering, along with the degree of cell proliferation and invasiveness in *KP200c* organoids, were proportional to the ratio of fibroblasts to cancer cells (Sup Fig. S3C). Consistent with these findings, re-expression of *mir-200c/141* in *KP200c* cancer cells co-cultured with lung fibroblasts reverted this invasive phenotype, restoring a spherical organoid morphology (Sup Fig. S3D, S3F). Hence, *miR-200* miRNAs act in lung cancer cells to restrict the metastatic phenotype, at least in part, by regulating the functional interaction between cancer cells and fibroblasts.

**Figure 3.**
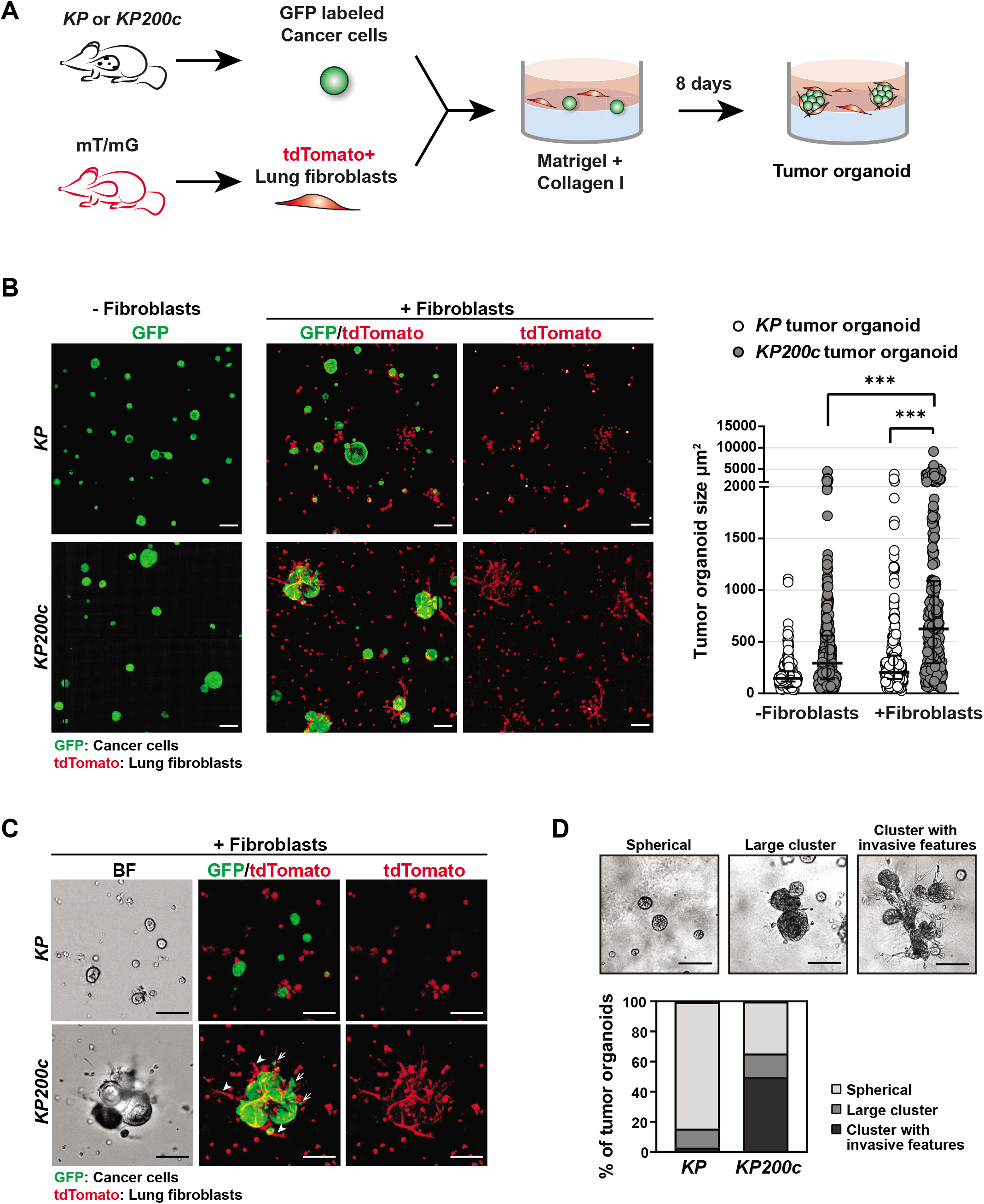
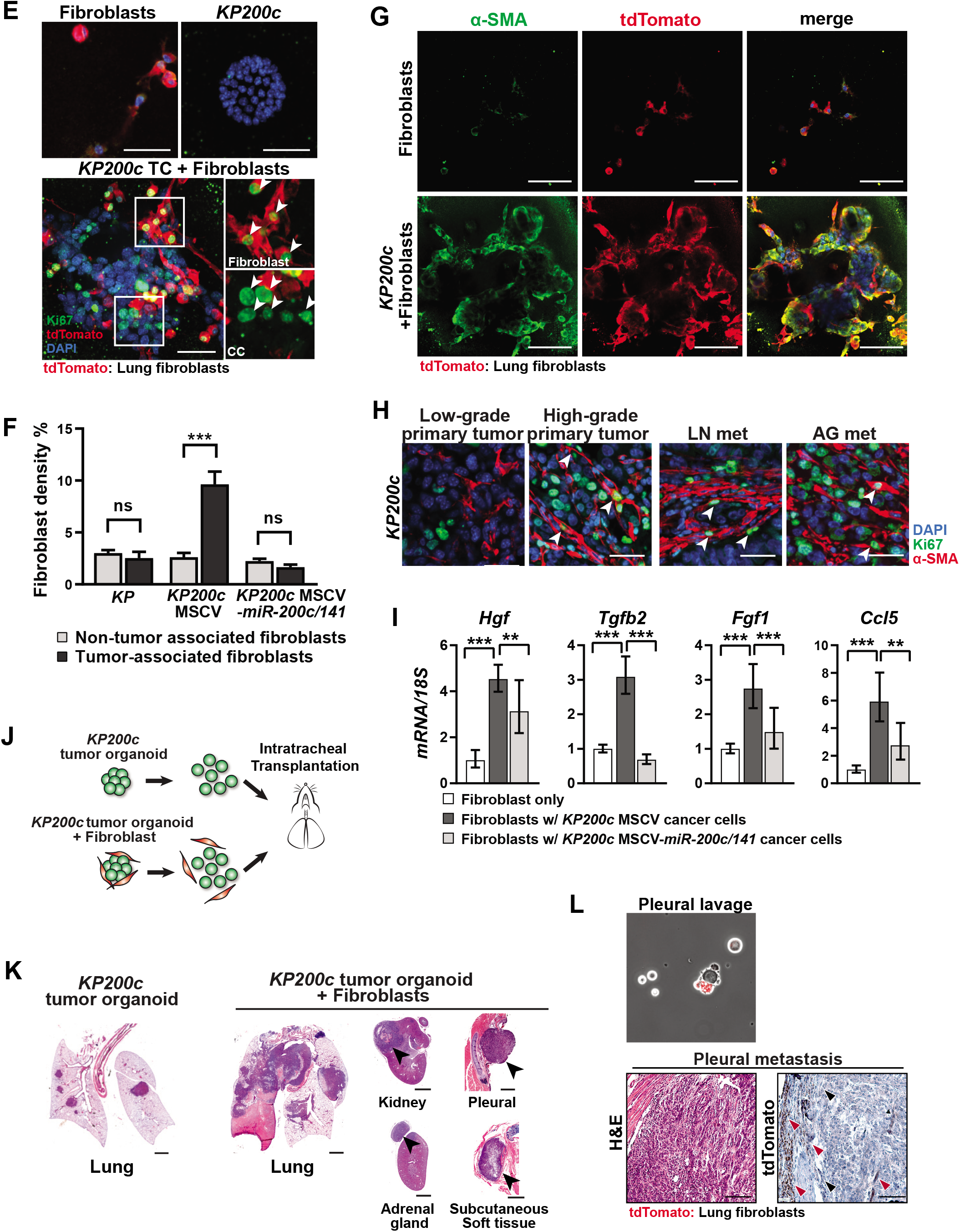
*miR-200* deficiency in cancer cells promote CAF activation to drive tumor metastasis. **A.** A diagram illustrating the tumor-fibroblast organoid co-culture model. GFP labeled *KP* or *KP200c* lung cancer cells are co-cultured with tdTomato labeled normal lung fibroblasts to form tumor organoids. **B, C, D.** *miR-200* deficiency in *KP* cancer cells induces metastatic features upon co-culture with lung fibroblasts. **B.** Fluorescence images (left) and quantitation (right) of tumor-fibroblast organoids indicate that *miR-200* deficiency in *KP* cancer cells, combined with fibroblast co-culture, induced tumor clustering and promoted metastatic cellular features. Scale bar, 100μm. *KP* alone: n=283, *KP+*Fibroblasts: n=237, *KP200c* alone: n=317, *KP200c+*Fibroblasts: n=256. *KP200c* alone vs. *KP200c*+Fibroblasts, *** *P*<0.0001, t=7.404, df=568; *KP200c*+Fibroblasts vs. *KP* +Fibroblasts, *** *P*<0.0001, t=7.894, df=488; unpaired two-tailed Student’s t-test. **C, D.** Representative images and quantification are shown for clustering and invasiveness in *KP* and *KP200c* tumor organoids with fibroblast co-culture. Scale bars, 100μm. **C**. *KP200c* organoids exhibit invasive features in cancer cells (arrows) and elongated morphology in co-cultured fibroblasts (arrowheads). **D**. Tumor organoids exhibit three morphologies: spherical organoids, large clusters, and large clusters with invasive cancer cell features. Analysis of 87 *KP* and 112 *KP200c* tumor organoids indicate a strong increase in invasive features in *KP200c* tumor organoids. **E.** Ki67 immunostaining in *KP200c* organoid with fibroblast co-culture indicates strong cell proliferation in both cancer cells and fibroblasts (tdTomato+). Scale bar, 100μm; white arrowheads, Ki67+ cells. **F.** *miR-200* re-expression in *KP200c* cancer cells decreases cell proliferation of tumor-associated lung fibroblasts in organoid co-culture, but fails to affect non-tumor associated fibroblasts. Density of tumor-associated fibroblasts and non-tumor-associated fibroblasts were quantified by measuring the tdTomato signal coverage. Error bar, s.d.. *KP* non-tumor-associated fibroblasts vs. tumor-associated fibroblasts, ns *P*=0.3823, t=1.021, df=3; *KP200c*-MSCV non-tumor-associated fibroblasts vs. tumor-associated fibroblasts, *** *P*=0.0006, t=6.484, df=6; *KP200c-MSCV-miR-200c/141* non-tumor-associated fibroblasts vs. tumor-associated fibroblasts, ns *P*=0.2456, t=1.440, df=3, unpaired two-tailed Student’s t-test. **G.** Co-culture of *KP200c* cancer cells and lung fibroblasts in organoids promotes α-SMA induction in fibroblasts. Scale bars, 100μm. **H.** Increased α-SMA+ CAF proliferation is observed in *KP200c* high-grade primary tumors and metastatic tumors, but not in *KP200c* MSCV low-grade primary tumors. Low-grade primary tumors, high-grade primary tumors, lymph node metastases, and adrenal gland metastases from *KP200c* mice were immunostained for Ki67 and α-SMA. White arrowheads, proliferating CAFs (Ki-67+, α-SMA+). Scale bars, 20μm. **I.** *miR-200* expression in *KP200c* cancer cells regulates the induction of specific cytokine and growth factors by CAFs. Lung fibroblasts co-cultured with or without control *KP200c* organoids or *miR-200* re-expressing *KP200c* organoids were isolated by flow cytometry, and *Hgf, Tgfb2, Fgf1* and *Ccl5* levels were quantified by real time PCR. Control *KP200c* cancer cells enhanced fibroblast production of pro-metastatic cytokines and growth factors, while *miR-200* re-expression in *KP200c* cancer cells reversed this phenotype. Error bar, s.d. All statistical analyses are performed using unpaired, two-tailed Student’s t-test. **J. K.** *KP200c* cancer cells primed with lung fibroblasts yield highly metastatic lung tumors in an orthotropic allograft tumor model 4 weeks after tumor transplantation. **J.** A cartoon illustrating the orthotropic allograft model to study lung cancer metastasis using co-cultured *KP200c* cancer cells and lung fibroblasts. *KP200c* tumor organoids without or with tdTomato+ lung fibroblast co-culture at 1:5 ratio for 5 days were dissociated, and single cell suspension containing 1×10^5^ cancer cells were subsequently transplanted to immune deficient *nu/nu* recipient mice through intratracheal injection. **K.** H&E staining reveals a highly metastatic tumor phenotype in primary tumors, lymph node metastases, and distant metastases generated from *KP200c* cancer cells primed by lung fibroblast co-culture. Scale bar, 500μm, black arrowheads, metastases **L**. Transplanted exogenous fibroblasts and endogenous fibroblasts from recipient mice both contribute to CAFs in metastatic tumor microenvironment. Disseminating tumor clusters from pleural cavity and pleural metastasis both contain transplanted tdTomato+ CAFs and endogenous tdTomato-CAFs. Scale bar=100μm, red arrowheads, tdTomato+ CAFs, black arrowheads, tdTomato-CAFs.

### *miR-200* deficiency in tumors promotes the proliferation and activation of CAFs

The interaction between *KP200c* cancer cells and stromal fibroblasts induced invasive features in neoplastic cells, while promoting their proliferation in 3D organoid culture (Fig 3E), but not in 2D culture (Sup Fig S3E). Increased fibroblast proliferation was only observed in the vicinity of *KP200c* tumor organoids (Fig. 3B), as tumor-associated fibroblasts exhibited a 3.8-fold increase in cell density compared to non-tumor-associated fibroblasts (Fig. 3F). No proliferation difference was observed between *KP* tumor-associated and non-tumor-associated fibroblasts (Fig. 3F). *KP200c* tumor-associated fibroblasts exhibited elongated morphology, induction of α-SMA, and infiltration into tumor organoids to form direct cell-cell contact with cancer cells (Fig. 3G). Consistently, highly proliferative α-SMA+ CAFs with an elongated morphology were prevalent in *KP200c* high-grade primary tumors, lymph node, and distant metastases, but not in low-grade primary tumors (Fig. 3H, Sup Fig S3G).

*miR-200* deficiency in lung cancer cells not only promotes the proliferation of surrounding lung fibroblasts, but also regulates their activity. A major function of CAFs is to secrete cytokines, chemokines, and growth factors to promote tumor growth, survival, and invasion^18^. We isolated tdTomato+ lung fibroblasts that were cultured alone or co-cultured with *KP200c* tumor organoids, and compared their expression of CAF-derived secreted factors. Fibroblasts cocultured with *KP200c* tumor organoids exhibited a strong induction of hepatocyte growth factor (HGF), TGF-β, fibroblast growth factor-1 (Fgf1) and Ccl5 (Fig. 3I), all of which have been reported to contribute to metastatic potential in different cancer models including lung adenocarcinoma^27–33^. Interestingly*, mir-200c/141* re-expression in *KP200c* tumor organoids reduced the induction of these growth factors and cytokines in co-cultured fibroblasts (Fig. 3I), suggesting that *miR-200* miRNAs in cancer cells regulate CAF gene expression via a cell nonautonomous mechanism. Moreover, treating *KP200c* cancer cells with HGF, TGF-β, or a combination of both is sufficient to induce the invasive behavior, resulting in enlarged invasive tumor organoid clusters (Sup Fig S3H).

### Interplay between *KP200c* cancer cells and fibroblasts promotes metastatic potential *in vivo*

Our findings implicate a complex interplay between cancer cells and CAFs regulated by *miR-200* miRNAs. To investigate the importance of cancer-fibroblast interactions in promoting metastases, we established an orthotropic allograft lung tumor model, in which we intratracheally transplanted lung cancer cells with or without primary lung fibroblasts into immune deficient recipient mice (Fig. 3J). Among two *KP* and three *KP200c* lung cancer lines tested, *KP* lines rarely grafted to form tumors, yet all *KP200c* lines successfully developed allograft lung tumors by 7 weeks (Sup Fig. S3I). When *KP200c* cancer cells were injected with co-cultured, tdTomato-labeled, primary lung fibroblasts, tumor growth was significantly accelerated with an increased tumor burden (Fig. 3K, Sup Fig. S3J). More significantly, within a short latency of 40 days, all mice exhibited early onset distant metastases affecting the pleural membrane in rib cage, pericardium, adrenal gland, kidney, and subcutaneous soft tissue (Fig. 3K, Sup Fig. S3K). In comparison, mice injected with *KP200c* cancer cells alone, while developing lung tumors, rarely developed metastases at this time point (Fig. 3K, Sup Fig. S3K). These allograft *KP200c* tumors likely gain metastatic potential eventually, as they promoted the proliferation and activation of endogenous stromal fibroblasts in recipient mice. Altogether, our data demonstrate that boosting cancer-fibroblast interactions in *KP200c* cancer cells significantly accelerated tumor outgrowth and metastases *in vivo*.

Surprisingly, when *KP200c* cancer cells were co-transplanted with primary lung fibroblasts into recipient mice, tdTomato+ fibroblasts could be detected within the disseminating cancer cell clusters in the pleural cavity and in distant metastases (Fig. 3L). Similarly, in *KP200c* mice, we observed cancer cell clusters that contained α-SMA+ CAFs inside blood/lymphatic vessels in the lung (Sup Fig S3L). Consistently, in a human patient with minimally invasive lung adenocarcinomas and lymph node micro-metastases, we detected clusters of cells containing α-SMA+ CAFs adjacent to CK19+ cancer cells in blood (Sup Fig. S3M), and observed lymph node micro-metastases encased by a single layer of α-SMA+ CAFs in the vicinity of lymphatic vessels (Sup. Fig S3N). These findings suggest that α-SMA+ CAFs can adhere to metastatic cancer cells during dissemination and extravasation, and prolonged tumor-CAF interaction could play a role throughout different stages of tumor metastasis.

### *miR-200* miRNAs repress *Jag1/Jag2* and inhibit Notch signaling in CAFs to repress metastasis

Cancer cells can convert lung fibroblasts into tumor-promoting CAFs via paracrine signaling or direct cell-cell interactions. Interestingly, *KP200c* cancer cells induced lung fibroblast proliferation and activation in a tumor organoid co-culture system that permitted cell-cell contact, but not in trans-well assays that separated cancer cells and fibroblasts in different compartments to prohibit cell-cell contacts (Fig. 4A). Hence, *miR-200* miRNAs likely regulate pathways for direct cell-cell interactions between cancer cells and CAFs.

**Figure 4.**
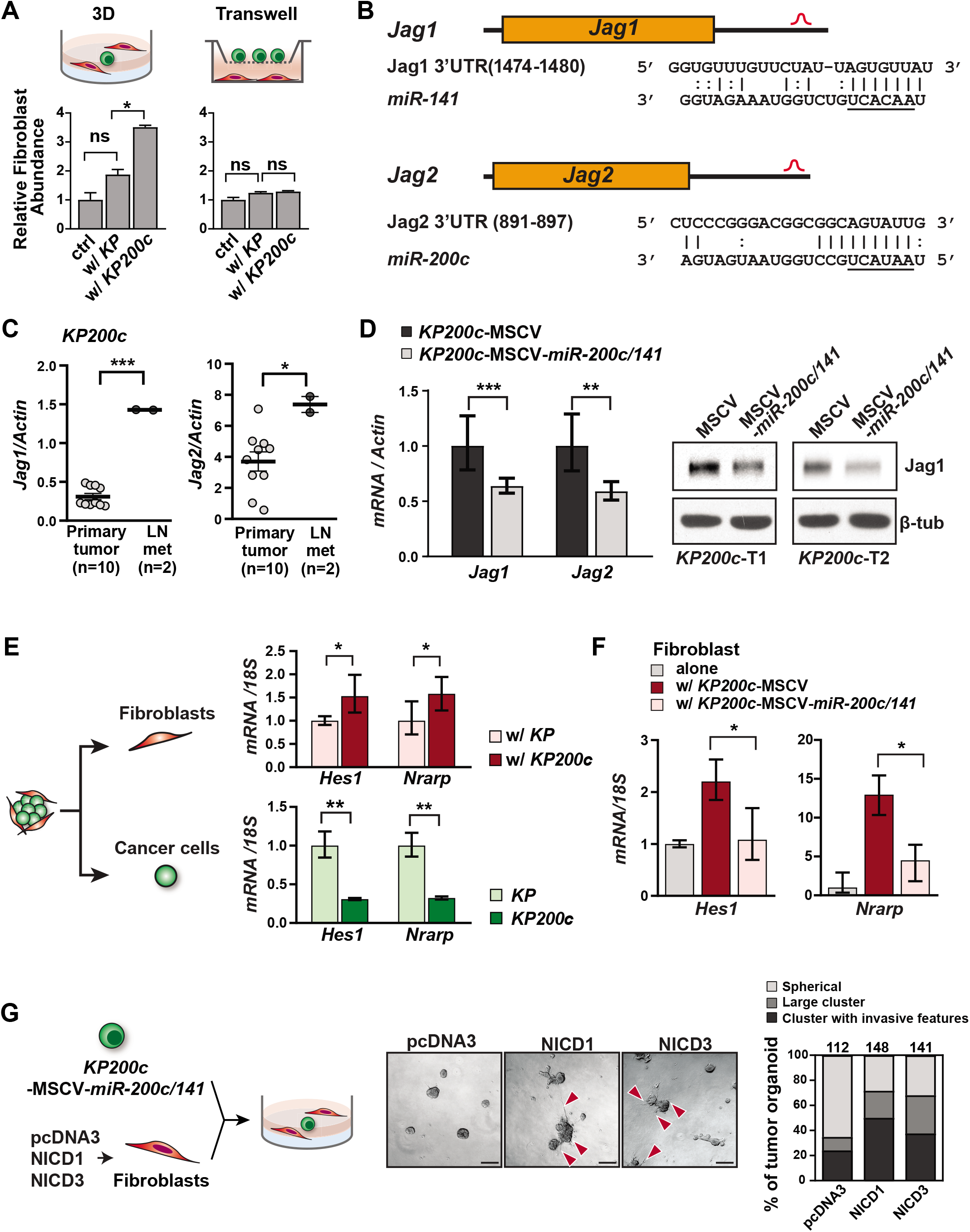
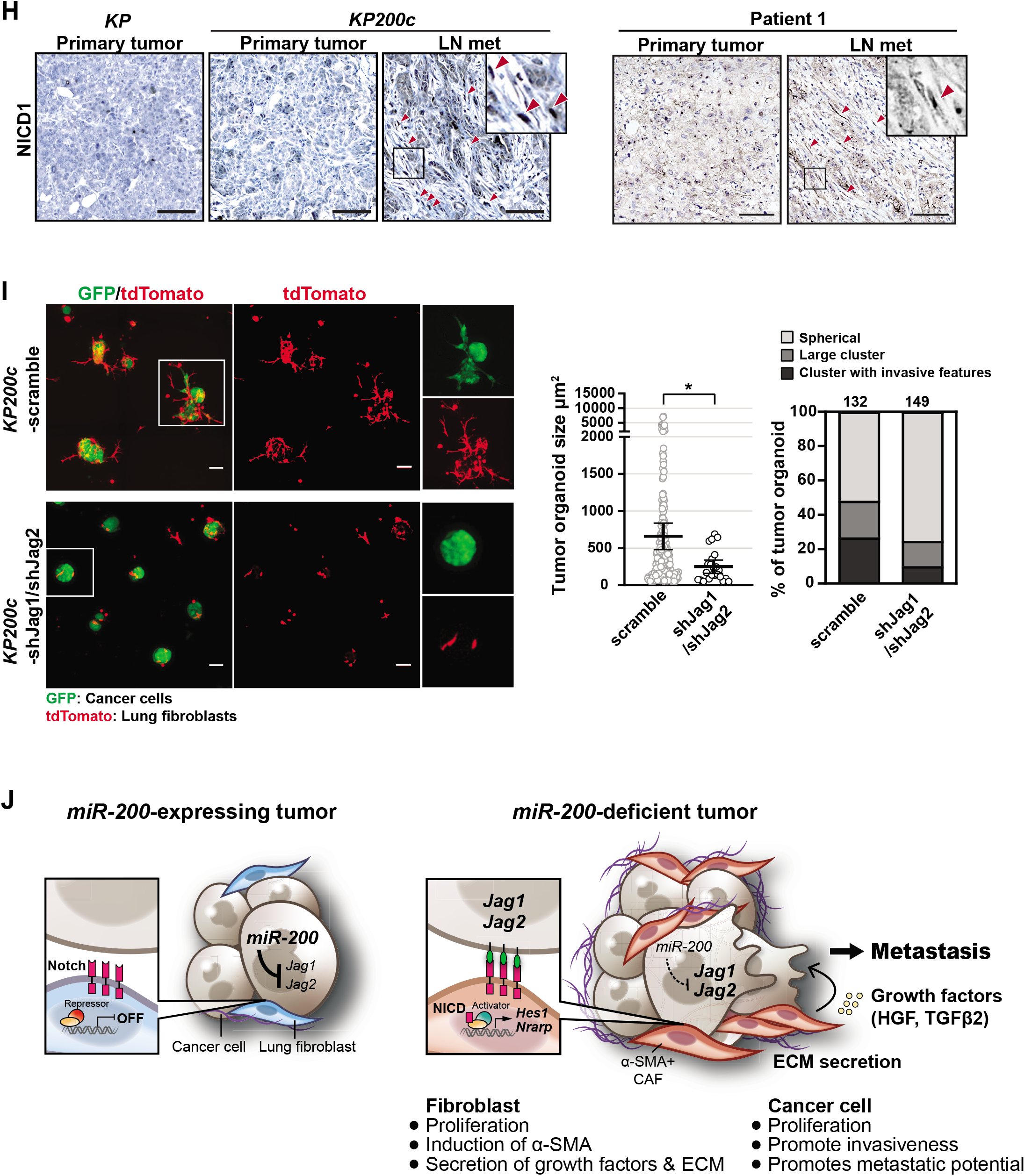
*miR-200* represses *Jag1/Jag2* in cancer cells and inhibits Notch signaling in CAFs. **A.** Direct cell-cell contact mediates the activation of co-cultured fibroblasts by *KP200c* cancer cells. Fibroblasts and *KP200c* cancer cells were co-cultured either in the organoid coculture or in transwell assays, and the abundance of fibroblasts was measured either by the percentage of tdTomato+ signal (in organoid co-culture) or by OD595 of crystal violet staining (in transwell assay). *KP200c* cancer cells promote proliferation of fibroblasts only in tumor organoid culture, where direct cell-cell contact was permitted. Error bars, s.d., Fibroblasts + KP tumor organoids vs. Fibroblasts + *KP200c* tumor organoids \**P*=0.0104, t=3.185, df=2, unpaired two-tailed Student’s t-test. **B.** Candidate *miR-141* and *miR-200c* binding sites in the 3’ UTR of *Jag1* and *Jag2*, respectively, as predicted by TargetScan. **C.** Metastatic *KP200c* tumors exhibit an induction of *Jag1* and *Jag2* compared to *KP200c* primary tumors. 10 *KP200c* primary tumors and 2 lymph node metastases were subjected to real time PCR for *Jag1* and *Jag2*. Error bars, standard error. *Jag1 KP200c* primary tumor vs. LN met *** *P*<0.0001, t=12.03, df=10, *Jag2 KP200c* primary tumor vs. LN met \**P=*0.0290, t=2.548, df=10, unpaired Student’s t-test. **D.** Overexpression of *mir-200c/141* in *KP200c* cancer cells suppresses the mRNA (left) and protein (right) expression of both *Jag1* and *Jag2*. Error bars, s.d.. **E.** Co-culture of lung fibroblasts and *KP200c* cancer cells induces Notch signaling in fibroblasts, but represses Notch signaling in cancer cells. tdTomato+ lung fibroblasts and GFP+ cancer cells were co-cultured as tumor organoids for 5 days, before each population was isolated using flow cytometry for real time PCR. Fibroblasts co-cultured with *KP200c* cancer cells exhibit an increase in Notch target genes (*Hes1* and *Nrarp*) compared to those co-cultured with *KP* cancer cells. When co-cultured with fibroblasts, *KP200c* cancer cells exhibit a decrease in Notch target genes compared to *KP* cancer cells. Error bar, s.d.. **F.** *miR-200* re-expression in *KP200c* cancer cells represses Notch targets *Hes1* and *Nrarp* in co-cultured fibroblasts. Error bars, s.d.. All statistical analyses were performed using unpaired, two-tailed Student’s t-test. **G.** Activated Notch signaling in fibroblasts promotes metastatic behavior of co-cultured, *miR-200* expressing cancer cells. Lung fibroblasts expressing mouse NICD1, NICD3, or control pcDNA3 vector were co-cultured with *mir-200c/141* expressing *KP200c* cancer cells, and invasive features of cancer cells were examined by light microscopy. Fibroblasts with activated Notch signaling, but not control fibroblasts, conferred a metastatic phenotype in *miR-200* expressing *KP200c* cancer cells in organoid culture. Scale bars, 100μm; red arrows, invasive cellular structures in cancer cells co-cultured with fibroblasts with activated Notch signaling. Morphology of tumor organoids was quantified as described in Fig. 3D. **H.** Metastatic *KP200c* mouse tumors and metastatic human lung adenocarcinomas both contain CAFs with strong expression of nuclear Notch1, compared to paired primary tumors. Scale bars, 100μm. Red arrows point to the CAF stained positive for activated Notch1. **I.** Knockdown of both *Jag1* and *Jag2* in *KP200c* cancer cells inhibits invasiveness in cancer cells and proliferation in fibroblasts when cultured as tumor organoids with lung fibroblasts. GFP labeled *KP200c* cancer cells expressing a control vector or shRNAs against both *Jag1* and *Jag2* were co-cultured with tdTomato labeled lung fibroblasts. Invasive features of cancer cells and cell proliferation of fibroblasts were analyzed by fluorescence microscopy (left). *Jag1* and *Jag2* double knockdown in *KP200c* cancer cells, when co-cultured with lung fibroblasts, restores a spherical morphology of tumor organoids (left), reduces the size of tumor organoids (middle), and decreases the invasive features of tumor organoids (right). Scale bar, 100μm. **J.** A model illustrating the role of *miR-200* miRNAs in lung cancer metastasis by regulating the interaction between cancer cells and fibroblasts. In *miR-200-*expressing cancer cells, *miR-200* targets *Jag1* and *Jag2;* hence, the cancer cells fail to activate Notch signaling in adjacent fibroblasts and lack metastatic potential. *miR-200* deficiency in cancer cells de-represses *Jag1* and *Jag2*, triggering Notch activation in neighboring CAFs, which promotes fibroblast proliferation and activation, ultimately enhancing the metastatic potential of the cancer cells.

Many studies suggest that a key function of *miR-200* miRNAs is to suppress EMT by targeting *Zeb1* and *Zeb2*^34–38^. However, primary and metastatic *KP200c* tumors remained largely intact in epithelial morphology (Sup Fig. S4A), and failed to exhibit *miR-200*-dependent regulation of Zeb1, Zeb2, Vimentin, and E-cadherin (Sup Fig. S4B, S4C). Furthermore, *Zeb1* and *Zeb2* knockdown in *KP200c* lung cancer cells did not alter the invasive morphology of *KP200c* tumor organoids (Sup Fig S4D, S4E). We cannot exclude the possibility that transient EMT still occurs during lung cancer metastasis *in vivo*, since *miR-200*-deficient lung cancer cells are more susceptible to TGF-β-induced EMT^34^. Yet our data strongly suggest that the *miR-200-EMT* pathway is not required for cancer-CAF interactions which play an essential role in promoting metastasis.

Gene ontology analysis on predicted *miR-200* targets revealed multiple candidates involved in cell-cell communication, including *Jagged1* (*Jag1*), *Jagged2* (*Jag2*), *Postn, Gjc1*, and *Pcdh9*, as predicted by RNA22 and TargetScan algorithms^39,40^. Notch ligands *Jag1 and Jag2* emerged as strong candidate *miR-200* targets, each containing a predicted *miR-141* or *miR-200c* binding site, respectively (Fig. 4B). Furthermore, their mRNA expression was elevated in *miR-200-* deficient metastatic cancer cells where lost the expression of both mir-200 loci (Fig. 4C, Sup Fig. S1D), but reduced upon *miR-200* re-expression (Fig. 4D). Hence, the di-cistronic miRNA cluster *mir-200c/141* acts coordinately to downregulate both *Jag1* and *Jag2* expression in cancer cells.

Notch signaling operates in diverse developmental and pathological processes. Notch ligands, including Jag1, Jag2, Dll1, and Dll4, bind to membrane-bound Notch receptors in neighboring cells, inducing a cascade of proteolytic events that ultimately release the Notch intracellular domain (NICD) which translocates into the nucleus^41–43^. In the nucleus, NICD interacts with DNA binding protein RBPJ and co-activator MAML to stimulate the transcription of Notch target genes, such as Hes, Hey, and Nrarp^44,45^. Notch expressing cells are often activated by cell surface ligands in adjacent cells, and Notch ligand producing cells are known to dampen their own Notch activity through cis-inhibition^46^. Consistently, *miR-200* deficiency in *KP200c* cancer cells resulted in elevated expression of Notch target genes *Hes1* and *Nrarp* in co-cultured fibroblasts, but a decrease in Notch target gene expression in *KP200c* cancer cells (Fig. 4E). This induction of Notch activity in co-cultured fibroblasts was reversed by re-expressing *miR-200c/141* in *KP200c* cancer cells (Fig. 4F).

To investigate the sufficiency of elevated fibroblast Notch signaling to promote the metastatic potential of cancer cells, we manipulated Notch signaling in lung fibroblasts and examined their effects on co-cultured tumor organoids. Since Notch1 and Notch3 are the main Notch receptors expressed in lung fibroblasts (data not shown), we activated Notch signaling in lung fibroblasts by overexpressing Notch1 NICD (NICD1) or Notch3 NICD (NICD3), and co-cultured them with *miR-200c/141-re-expressing KP200c* tumor organoids. While *miR-200* expression abolished metastatic features of *KP200c* tumor organoids when co-cultured with normal lung fibroblasts (Sup Fig. S3F), co-culture with NICD1 and NICD3 expressing fibroblasts reversed the effects of *miR-200* re-expression in *KP200c* tumor organoids, promoting cancer cell clustering and invasive cellular features (Fig. 4G). In line with these findings, CAFs in metastatic *KP200c* tumors exhibited specific activated Notch1 nuclear staining (Fig. 4H, left), indicative of a strong Notch activation in these cells. Similar observations were made in two human lung adenocarcinoma samples, where activated Notch1 nuclear staining was evident in CAFs associated with metastases (Fig. 4H, Sup Fig. S4G).

Consistent with *Jag1* and *Jag2* being the key *miR-200* targets that mediate tumor-CAF interactions, knocking down *Jag1* and *Jag2* in *KP200c* cancer cells restored a spherical morphology in tumor organoids, abolished tumor clustering and invasive features, and reduced fibroblasts elongation and expansion (Fig. 4I). Hence, *Jag1* and *Jag2* derepression caused by *miR-200* loss in cancer cells triggered Notch activation in adjacent CAFs, which promoted fibroblast proliferation and activation, ultimately enhancing the metastatic potential of cancer cells (Fig. 4J).

## Discussion

*miR-200* miRNAs act as potent suppressors of lung cancer metastasis, and complete *miR-200* inactivation was observed in metastatic lung adenocarcinomas in both *KP* and *KP200c* models. The *KP200c* model is among the best for studying lung adenocarcinoma metastasis, owing to its short latency and faithful recapitulation of patient pathology. Metastatic *KP200c* lung adenocarcinomas faithfully recapitulate the desmoplastic stroma found in human metastatic tumors^9,47^, characterized by a pro-metastatic niche filled with SMA+ CAFs. These SMA+ CAFs, derived from normal lung fibroblasts, strongly foster the metastatic potential of cancer cells, at least in part, by secreting cytokines and growth factors (Fig. 3G, 3I). A recent study showed that human metastatic lung adenocarcinomas possess transcriptomes encompassing a developmental continuum ranging from stem cell-like cells to regenerative progenitors in lung epithelia^48^. It is likely that CAFs in the desmoplastic microenvironment not only promote tumor cell proliferation, but also create a pro-metastatic niche to induce developmental plasticity in cancer cells. This is consistent with the reports that Wnt-expressing fibroblasts establish a stem cell niche for AT2 cells in normal development and for cancer stem cells in lung adenocarcinoma^49,50^. Given the heterogeneity of the tumor microenvironment, CAF-dependent pro-metastatic niches are likely to require interactions between CAFs and other stromal cell types in lung adenocarcinomas.

In most studies, *miR-200* miRNAs are characterized as key inhibitors of EMT by repressing *Zeb1* and *Zeb2*^34–38^. Yet EMT may not be essential for metastasis in *KP200c* lung adenocarcinomas, as overt EMT was not observed in metastatic *KP200c* tumors, and *Zeb1* and *Zeb2* knockdown failed to reverse invasiveness in tumor organoids (Sup Fig. S4A, S4D, S4E). Rather, miR-200 deficiency in *KP200c* cancer cells leads to increased Jag1 and Jag2, elevating Notch signaling in neighboring fibroblasts to promote their conversion into α-SMA+ CAFs, thereby establishing a pro-metastatic niche. This Notch activation in CAFs is strongly associated with metastatic lung adenocarcinomas in both mouse models and human patients (Fig. 4H), highlighting the importance of the *miR-200*/Jag/Notch axis in orchestrating cancer-CAF interactions to promote metastasis in lung adenocarcinomas. Notch activation in lung fibroblasts also induces α-SMA expression and TGF-β secretion during lung fibrosis^51^. Hence, Notch signaling regulates fibroblast dependent, cell-cell interactions in multiple lung pathological processes.

The effects of Notch signaling are complex in various tumor types. Notch activation in fibroblasts promotes CAF activation and tumor progression in lung, breast, and prostate cancer^52,53^, and the loss of Notch signaling in CAFs promotes an oncogenic tumor microenvironment in squamous cell carcinomas^54,55^. It is important to recognize that the unique cellular heterogeneity found within each tumor microenvironment could foster a complex crosstalk between cancer cells and stromal cells. In both mouse and human metastatic lung adenocarcinomas, *miR-200* deficient lung cancer cells could also induce Notch signaling in other stromal cell types, which in turn promoted metastasis. Taken together, *miR-200* miRNAs regulate Notch signaling, which mediate important cell-cell interactions between cancer cells and stromal cells to suppress lung cancer metastasis.

## Acknowledgements

We thank H. Nolla, H. Kartoosh and A. Valeros from for FACS support; A.M. Li for establishing a collaboration with Dr. Bradley’s lab; A. Holly, J.Y. Lee and F. Ives from Molecular Imaging Center at UC Berkeley and M. West and P. He from HTS facility at UC Berkeley for imaging assistance; Functional Genomics Laboratory at UC Berkeley for genomics services, C. Hung for experimental help and proofreading the manuscript, S. Chen for revision advice, G. Lee, A. Nazarenko, T. Colston and S. Yue for experimental help and mouse colony management; C. Fellmann for providing SGEP and SGEN vectors; M. Junttila, R. Molina and J. Long from Genentech for intranasal injection protocol. H. Ji and F. Li for *Kras^G12D/+;^Lkb1^flox/flox^* tumor samples. N. Arpaia, K.A. Kaiser and all He lab members, in particular N. Okada, M. Bennett, E. Ho and J. Schwedler for suggestions and discussions. L.H. is a Thomas and Stacey Siebel Distinguished Chair Professor, who is supported by a Howard Hughes Medical Institute (HHMI) Faculty Scholar award, a Bakar Fellow award, at UC Berkeley, and several grants from the National Institutes of Health (NIH, R01CA139067 and 1R21OD027053). M.M. W. was supported by NIH R01-CA175336 and R01-CA204620. C.H.C was funded by an American Lung Association Fellowship. B. S. was supported by Grant No.81572269, National Natural Science Foundation of China.

## Author Contribution

B.X., B.S., M.M.W., and L.H. conceived, designed, and directed the study. B.X. performed most mouse experiments and generated the majority of the mouse data for this study; B.S. performed most experiments involving human samples, and has contributed his clinical expertise to this study; C.H.C and M.M.W. generated and isolated *KP* tumors and metastases for miRNA analysis. J.C. and A.B. performed survival analysis of human data from TCGA. J.E.W. performed pathology analysis on all the mice tumors. C.C., N.S., C.S.F. and M.K. performed mouse tumor histology, IHC staining and molecular cloning experiments. Y.W. and C.C. designed and provided miRNA TaqMan analysis. H.M.P and A.B. generated *miR-200c/141 null* mice. M.T.M. generated *mir-200c/141^lacZ^* mice. B.X. and L.H. wrote the manuscript with input from all authors.

## Data and materials availability

All data is available in the main text or the supplementary materials. All data and reagents will be available to any researcher for purposes of reproducing or extending the analysis.

## Material and Methods

### Animals

*Kras*^LSL-*G12D/+*^*;Trp53^flox/flox^;R26^LSL-tdTomato/+^* mice have been described previously^12,14,56^. *miR-200c/141* and *miR-200b/200a/429 null* mice were acquired from The Wellcome Trust Sanger Institute^16^. Mice of *Kras*^LSL-*G12D/+*^*;Trp53^flox/flox^* genotype were crossed with *mir-200c/141^-/-^* mice to generate *Kras*^LSL-*G12D/+*^*;Trp53^flox/flox^;mir-200c/141^-/-^* (*KP200c*) mice. *miR-200c/141^lacZ^* mice were acquired from MMRRC and Michael McManus lab^26^, they were crossed with mice expressing Flippase (ACTB:FLPe B6J, JAX) to generate *miR-200c/141^flox^* alleles. *KP200c* genotype were crossed with *mir-200c/141^flox/flox^* mice to generate *Kras*^LSL-*G12D/+*^*;Trp53^flox/flox^;mir-200c/141^flox/-^* (*KP200cCKO*) mice. Lung tumorigenesis was induced by intranasal instillation of 5×10^6^ PFU of recombinant Ad-Cre virus (University of Iowa Gene Transfer Vector Core) or by intratracheal intubation of 1×10^5^ PFU of Lenti-Cre in 8-week-old mice^13^.

### Human primary and metastatic tumor samples of lung adenocarcinoma

Human tumor and blood samples were collected from patients approved by the Institutional Review Board of Shanghai Pulmonary Hospital, School of Medicine, Tongji University. The eight pairs of matched primary and metastatic tumor samples were collected from patients who were diagnosed with clinical stage IIa/IIb/IIIa, pathologically confirmed lung adenocarcinoma and underwent pulmonary lobectomy and lymphadenectomy at Shanghai Pulmonary Hospital. The two liver metastases were obtained from palliative surgical resection of relapse lung adenocarcinomas. The samples of early-stage lung cancer were obtained from patients with stage Ia adenocarcinoma by lobectomy or segmentectomy. The patients’ tumor tissues were made into FFPE blocks and subjected to H&E staining or IHC routinely. The blood samples for isolating circulating tumor cell clusters were collected from patients with lung adenocarcinoma at stage IV.

### Isolation of mouse tumor cells and mouse lung fibroblast

tdTomato+ tumor cells for miRNA profiling were collected from *KPT* mice by flow sorting as described previously^56^.

Cell lines were derived from individual tumors from *KP* and *KP200c* mice from 4 months after tumor initiation, and created by collagenase I (20U/ml, Worthington Biochemical)/ collagenase IV (800U/ml, Worthington Biochemical) digestion and mechanical dissociation.

Normal lung fibroblast expressing tdTomato were collected from *ROSA^mT/mG^* mice^57^ using protocol described previously^58^ with modifications. Following lung harvest, the tissue was cut into small pieces and lightly digested with Liberase, and then cultured on scratched surface in Dulbecco’s modified Eagle’s medium (DMEM) supplemented with 10% fetal bovine serum (FBS), 100 units/ml penicillin, 100 μg/ml streptomycin and 0.25 μg/ml amphotericin B (serum-free medium). Fibroblasts are highly migratory and proliferative compared to other cell types, they will attach to scratching marks and form outgrowth after 1 week. The fibroblast will be expanded and used 3-7 passages following primary culture.

### microRNA assays

Total RNA was extracted from isolated tdTomato+ *KPT* tumor cells using TRIzol (Thermo Fisher Scientific) according to manufacturer’s instruction. For Taqman microRNA assay, cDNA was synthesized with preamplification from total RNA using Megaplex™ RT Primers, Rodent Pool B v3.0 (Applied Biosystems™, 4444292), Megaplex™ PreAmp Primers, Rodent Pool B v3.0 (Applied Biosystems™, 4444308), TaqMan^®^ MicroRNA Reverse Transcription Kit and TaqMan™ PreAmp Master Mix Kit according to manufacturer’s instruction. Subsequently, quantitative-PCR was carried out using TaqMan™ Array Rodent MicroRNA A+B Cards Set v3.0 (Applied Biosystems™, 4444909). The data were analyzed using ViiA 7 Software and R program. Individual microRNA and mRNA were analyzed as previously described^59^. The primers used here are shown in Supplemental Table S3.

### Gene expression analysis of the TCGA data set

LUAD data was downloaded from TCGA and normalized using upper quantile. All patients were into two groups - “lower” and “higher”, according to the expression of the mir-200 family. The “lower” group included patients for whom all five mir200 expression values were lower than or equal to the median expression value of the corresponding mir200s in the normal control samples. The “higher” group included the rest of patients.

Survival analysis was performed to compare these two groups, as shown in Figure 1E, using the package “survival” in R (https://cran.r-project.org/web/packages/survival/index.html). *P*-value was determined by Log rank test, to quantify the significance of the differences in survival time between the two patient groups.

### Three-dimensional tumor organoid culture for tumor-stromal interaction

Lung tumor cells and primary lung fibroblast were routinely cultured in DMEM supplemented with 10% FBS, 100 units/ml penicillin, 100 μg/ml streptomycin. The tumor organoid culture was performed using protocol described previously^60^ with modifications. 8-well chamber slides were coated with 100ul Matrigel (354234, Corning)/ Collagen I (0.5mg/ml, 354236, Corning) mixture. The slide was incubated at 37 °C for 30 minutes to allow the mixture to solidify. Tumor cells and fibroblast cells were trypsinized and resuspended in growth media with supplement at 2×desired density and mix with equal volume of 10% Matrigel/Collagen I mixture. The cell/gel mixture were immediately plated on top of the matrix coating and cultured at 37°C, 5% CO_2_. From day 2 to day 8, 3D structures were monitored for invasive structure.

### Orthotopic transplantation experiment

Tumor cells and fibroblast were co-cultured in suspension for 5 days prior to transplantation experiment. To prepare the suspension culture, the dissociated tumor cells and fibroblast are mixed at 1:5 ratio and seeded at 5×10^4^ cells/ml density in ultra-low attachment dishes (Corning, CLS3262) in DMEM/F12 supplemented with human recombinant EGF (20 ng/mL, BioLegend), bFGF (20 ng/mL, PeproTech), B27 (Gibco by ThermoFisher) and 4μg/mL Heparin (StemCell Technology). On the day of transplantation, tumor organoids were dissociated into single cells using trypsin/EDTA, and the percentage of tumor cells and tdTomato+ fibroblasts was determined by FACS. Single cell suspension containing 1×10^5^ tumor cells were resuspended in 50μl RPMI1640 with 10mM EDTA and injected into the lung of *nu/nu* mice (Taconic) by intratracheal intubation. The tumors were collected either at 40 days after the tumor injection or at their terminal stage. The University of California Berkeley Animal Care and Use Committee approved all animal studies and procedures.

### Histopathology and immunohistochemistry

For determining tumor histology, whole lungs were fixed overnight in 10% buffered formalin (Fisher Scientific, no. SF100-4), dehydrated in a graded series of ethanol solutions, then embedded in Paraffin Wax (ACROS Organics). The lung blocks were sectioned at 5μm thicknesses, mounted on glass slides, and stained with hematoxylin and eosin (H&E) using standard procedures. Lung and tumor areas were determined using Image Viewer (Ventana Medical Systems/Roche). Each tumor was given a score of grade 1–5^14^. Grades 1 and 2 were classified as low-grade tumors, and grades 3–5 were classified as high-grade tumors.

For IHC, paraffin sections were deparaffinized, dehydrated, and subjected to heat-induced antigen retrieval in a pressure cooker using Trilogy™ (Cell Marque, 920P). Slides were incubated for 10 min with 3% H_2_O_2_, blocked for 1h with 10% goat serum in phosphate-buffered saline (PBS) containing 0.05% Triton X-100, and incubated with primary antibody overnight in blocking buffer. Horseradish peroxidase (HRP)-conjugated secondary antibodies were incubated for 2 h at room temperature, with a 1:400 dilution in blocking buffer. For CC10, Nkx2.1 and p-Erk staining, a biotinylated horse anti-rabbit secondary antibody (Jackson ImmunoResearch, 711-065-152) was used at 1:400 as above, followed by incubation with Vectastain elite ABC reagent (Vector Laboratories, PK-7100) for 30 min. Peroxidase was then visualized by DAB staining (Vector Laboratories, SK-4100) and counterstaining with Gill’s hematoxylin solution.

### Detection of circulating tumor cell clusters

The blood for circulating tumor cell clusters detection was obtained from patients with stage IV lung adenocarcinoma. The previously described subtractive enrichment approach^61^ was modified to detect circulating tumor cell clusters. Briefly, 5 mL heparin-anticoagulated blood was added with 10 volume of Tris-buffered NH4Cl for erythrocyte lysis, and centrifuged at 500g for 10 min. The pellets were resuspended in 0.2 mL PBS and re-lysed with 1 mL OptiLyse C (Beckman Coulter) for 15 min. After centrifugation, the pellets were dispersed and fixed in 1% paraformaldehyde PBS for 10 min, washed and resuspended in 0.5 mL 0.1% BSA PBS. 0.5 mL CD45 Dynabeads (ThermoFisher scientific) was added to deplete leukocyte fraction according to the manufactory’s instruction. The CD45-depleted fraction was smeared on 3 positive charged slides. Cell clusters were observed and labeled on microscope for the following H&E or immunofluorescent staining.

### Antibodies

The following monoclonal (mAb) and polyclonal (pAb) primary antibody were used for immunoblotting, IF and IHC: Prosurfactant protein (SPC) pAb (Millipore, AB3786), Uteroglobin (CC10) pAB (Millipore, ABS1673), pErk1/2 pAb (Cell Signaling Technology, 4370), TTF1 (Nkx2.1) pAb (Abcam, ab76013), activated Notch pAb (Abcam, ab8925), Jag1 mAb (Cell Signaling Technology, 70109), α-SMA mAb (Invitrogen, 14-9760-82), Ki67 mAb (Cell Signaling Technology, 12202), RFP pAb (Rockland, 600-401-379).

### Jag1/Jag2 knockdown

The Jag1 knockdown lentiviral expression vector were constructed into pRRL-SFFV-GFP-miR-E-Puro (SGEP) vector and the Jag2 knockdown lentiviral expression vector were constructed into pRRL-SFFV-GFP-miR-E-Neo (SGEN) vector^62^. The oligo sequences encoding Jag1 and Jag2 shRNA were designed using splashRNA algorithm^63^ (Supplementary Table S4), and cloned into XhoI/EcoRI sites. The shRNA lentiviral vectors were package in HEK293 cells together with pVSV-G and pMD-2 as previously described. Five shRNAs were tested for each gene, and the ones with best knockdown effect were used to generated double knockdown cell line. Cells transduced with both shJag1 and shJag2 were selected 48 hr post infection using 2μg/ml puromycin (Gibco by ThermoFisher) and 500μg/ml Geneticin (Gibco by ThermoFisher).

## Supplementary figure legend

**Supplementary Figure S1.**
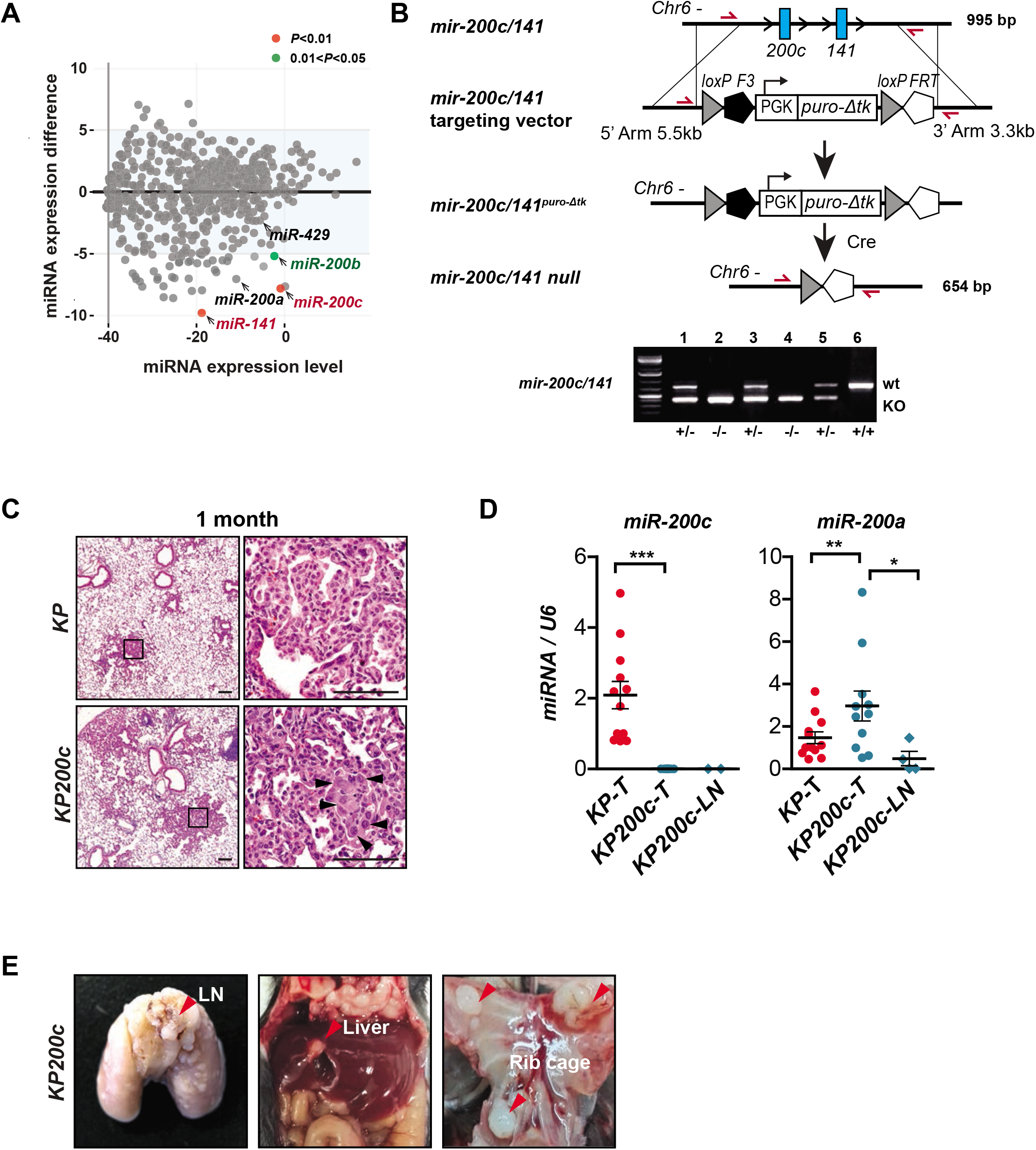
*miR-200* deficiency induces rapid metastasis in *KP* lung cancer model. **A.** *miR-200c* and *miR-141* are highly expressed in *KPT* primary tumors and significantly downregulated in metastases. A scatterplot shows the expression difference between the *KP* primary tumors and metastases (y axis, ΔCt_*KPT*.met_ – ΔCt_*KPT*.primary_) versus total miRNA expression level of *KPT* primary tumors and metastases (x axis, - [Δ_Ct*KPT*.primary_+Δ_Ct*KPT*.met_]). The *miR-200* family members are labeled with color corresponding with their *P* value range. **B.** Schematic illustration of *mir-200c/141* targeting and knockout strategy. Two homologous arms flanking the *mir-200c/141* locus mediate gene-specific targeting by homologous recombination. Targeting replaces *mir-200c/141* with a selection cassette containing a PGK-driven puromycin selection marker and a truncated TK variant (Δtk) flanked by loxP sites. Future puromycin selection and crossing with Cre transgenic mice leads to removal of reporter to generate *mir-200c/141* knockout mice. Red arrows indicate the position of the PCR primers used for genotyping. Bottom, homozygous deletion was confirmed by PCR. **C.** The appearance of pleomorphic nuclei in early stage *KP200c* primary tumor. Representative images of H&E staining of lung sections from *KP* and *KP200c* mice 1 month after Adeno-Cre infection are shown. Scale bar, 100μm. Arrows, cancer cells with pleomorphic nuclei. **D.** Loss of *miR-200* expression in the *KP200c* primary tumors. *miR-200c* and *miR-200a* expression was measured by real time PCR in the *KP* primary tumors (n=12), *KP200c* primary tumors (n=11) and *KP200c* LN metastases (n=4). Error bars, s.e.m. **E.** Occurrence of metastases in multiple sites in *KP200c* mice. Photos of LN metastasis, liver metastasis, and pleural metastases on the pleural membrane in *KP200c* mice are shown.

**Supplementary Figure S2.**
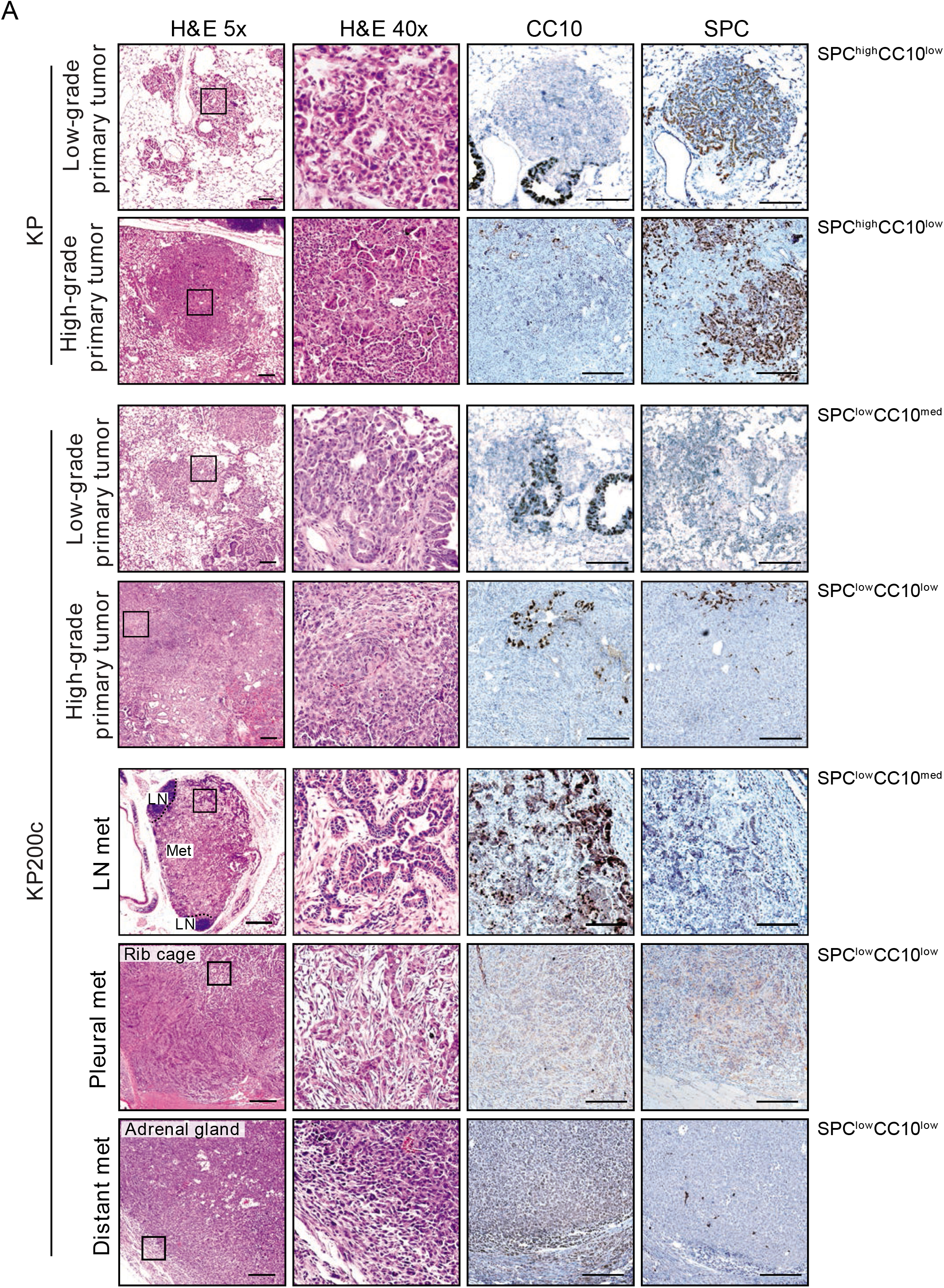
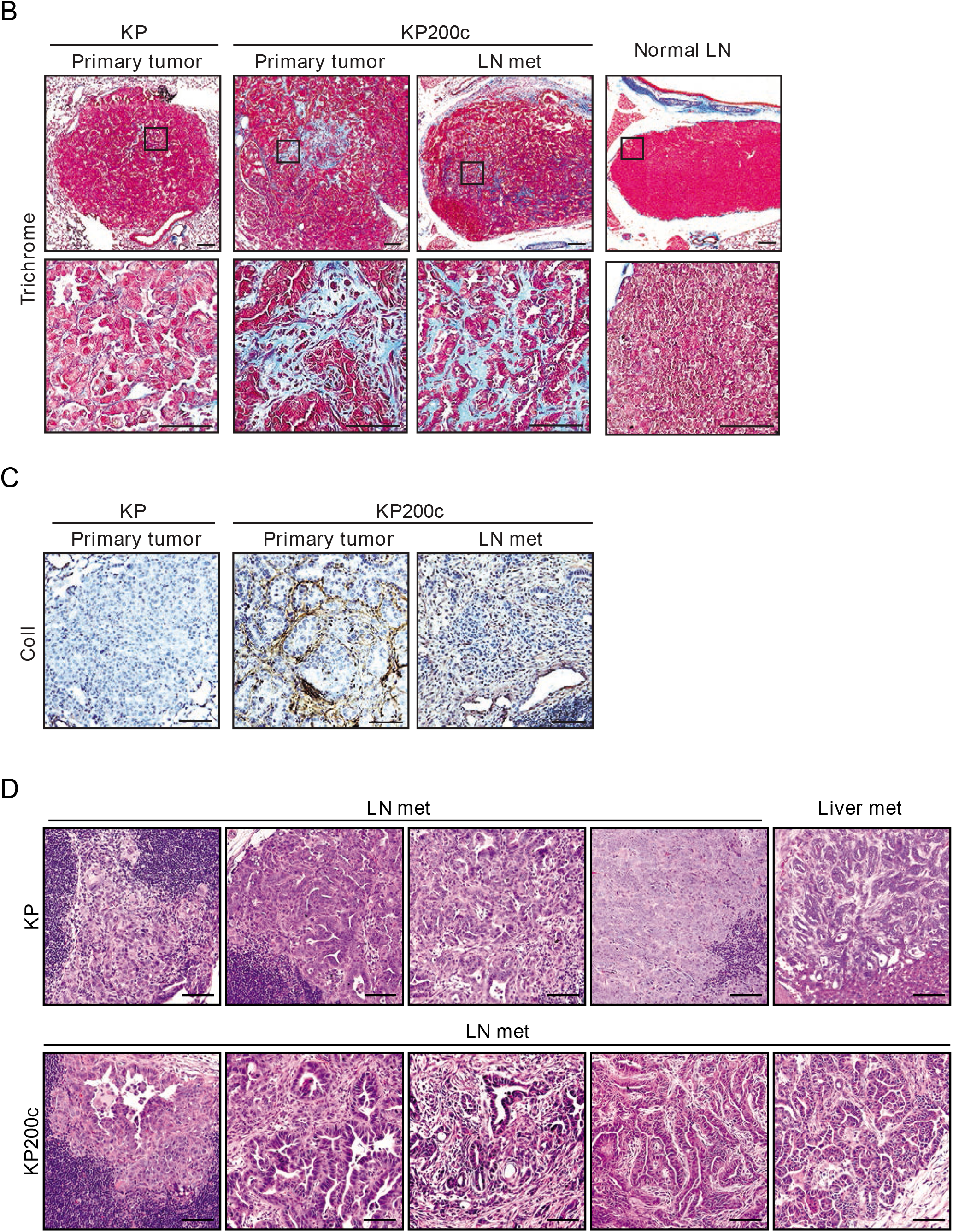
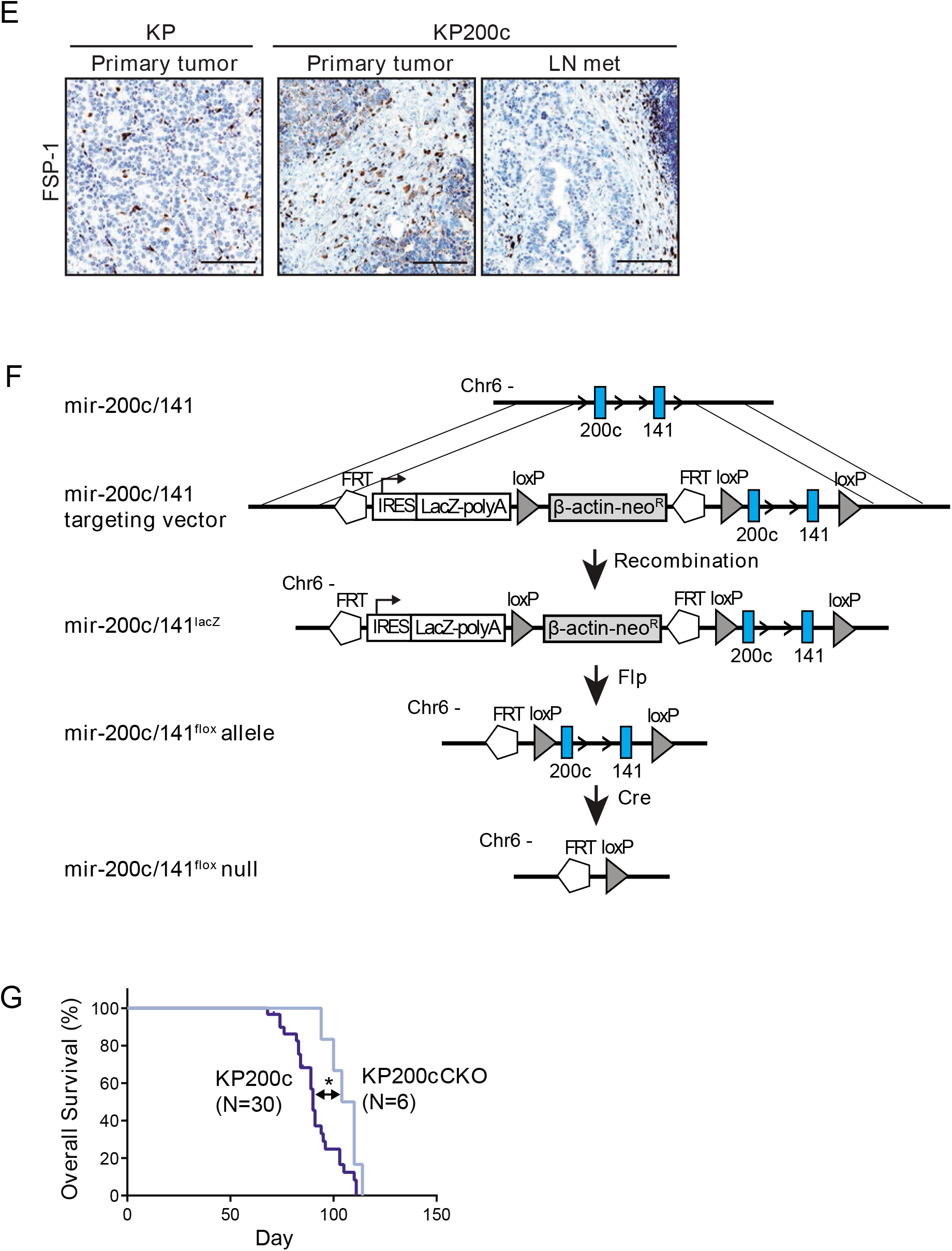
Characterization of desmoplasia phenotype in metastatic *KP200c* lung adenocarcinomas. **A.** Loss of cell lineage markers in metastatic *KP200c* tumors. H&E and IHC staining of *KP* and *KP200c* low-grade and high-grade primary tumors and *KP200c* LN metastasis, pleural metastasis, and distant metastasis with lung lineage markers including CC10 (a marker for Club cells) and SPC (a marker for alveolar type 2 cells) are shown. The SPC/CC10 status is indicated on the right. Scale bar, 100μm. **B.C.** Enrichment of collagen-rich extracellular matrix in metastatic *KP200c* tumors. Representative images of **B.** Masson’s trichrome staining and **C.** IHC staining for Collagen I of *KP* primary tumor, *KP200c* primary tumor, and *KP200c* metastasis are shown. In trichrome staining, collagenous fibers are stained blue. Normal lymph node (LN) from uninfected mouse is shown as control. Scale bar, 100μm. **D.** Lack of stromal desmoplasia in metastatic *KP* tumors. Representative images of H&E staining of *KP* LN metastases, liver metastasis and *KP200c* LN metastases are shown. Scale bar, 100μm. **E.** No change in FSP-1-expressing CAFs in metastatic *KP200c* tumors. Representative images of IHC staining for FSP-1 in *KP* primary tumor, *KP200c* primary tumor, and LN metastasis are shown. Scale bar, 100μm. **F.G**. *miR-200* deficiency in lung cancer cells promotes local and distant metastases. **F.** Schematic illustration of *mir-200c/141* targeting and generation of conditional knockout allele. Two homologous arms flanking the *mir-200c/141* locus mediate gene-specific targeting by homologous recombination. Targeting leads to insertion of a cassette containing LacZ-polyA flanked by FRT and *mir-141/200c* flanked by loxP sites. Future neomycin selection and crossing with flippase transgenic mice leads to removal of reporter to generate a conditional *mir-200c/141* knockout allele (*mir-200c/141^flox^*). Cre virus induces deletion of *mir-200c/141* in infected cells. **G.** Kaplan-Meier survival curve of *KP200c* and *KP200cCKO* mice after Adeno-Cre administration (5×10^6^ Ad-Cre PFU/mouse), animal number N is indicated in the figure, Log rank Test \**P*-value= 0.0409.

**Supplementary Figure S3.**
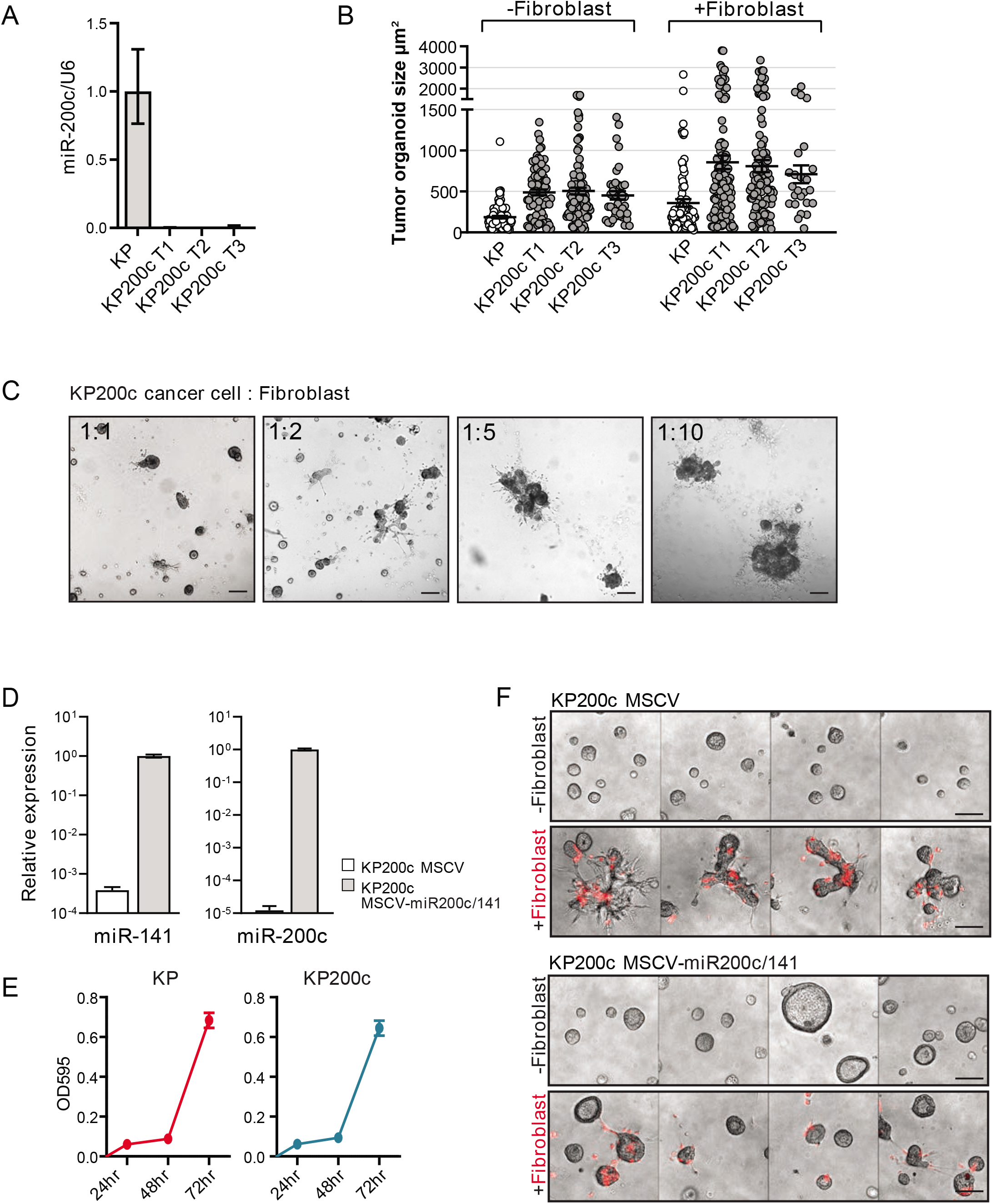
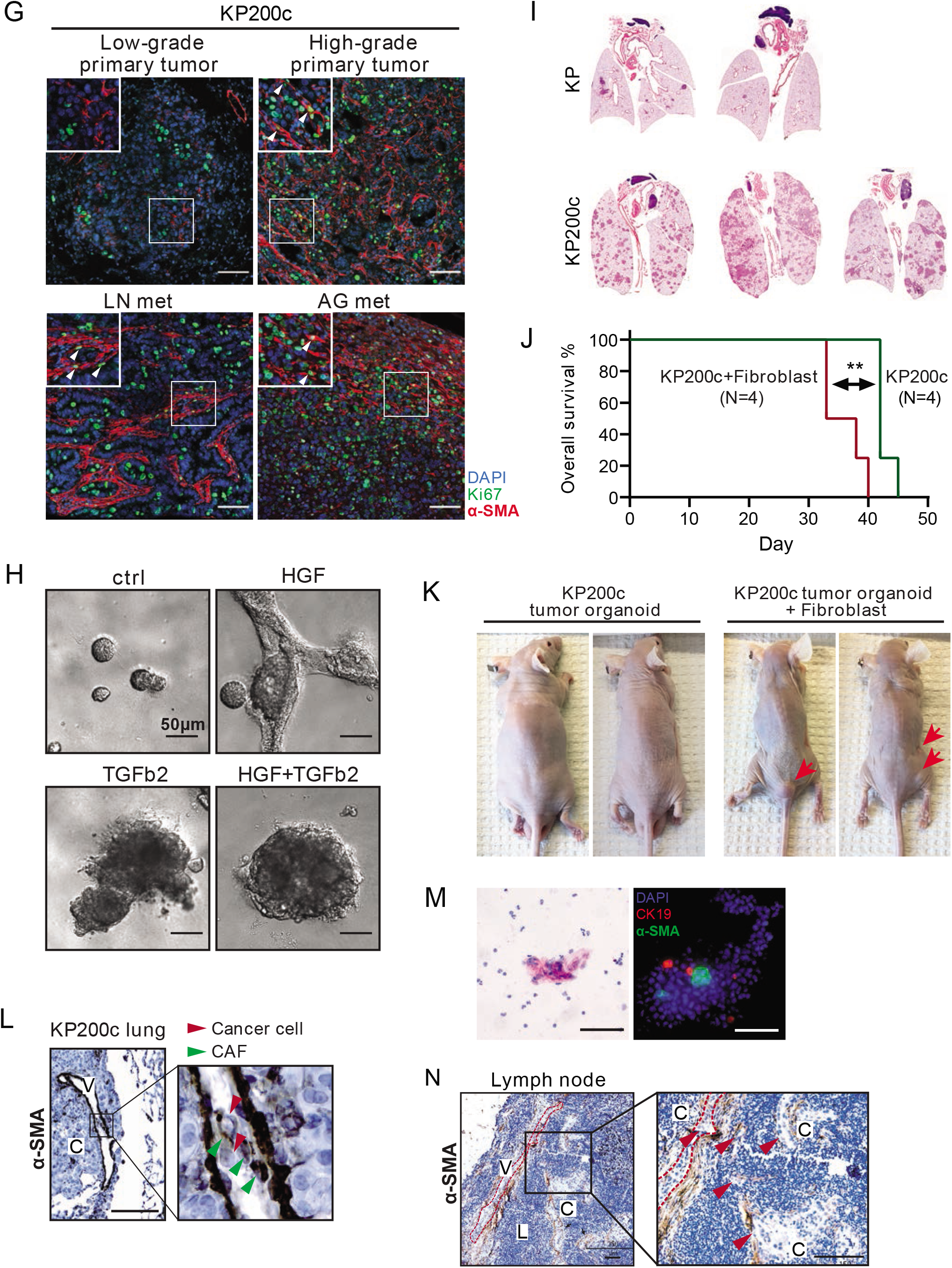
*miR-200* deficiency promotes CAF activation and metastatic features in cancer cells. **A.** Loss of miR-200 expression in *KP200c* cancer cell lines. Expression level of *miR-200c* in one *KP* and three *KP200c* cancer cell lines was quantified by real time PCR, and U6 was used as housekeeping control. Error bar, s.d. **B.** Increased size of *KP200c* tumor organoids upon co-cultured with lung fibroblasts in multiple *KP200c* lines. One *KP* and three *KP200c* cancer cell lines were co-cultured with or without lung fibroblasts, and tumorsphere size was measured by calculating the GFP (tumor) area coverage. More than 200 tumorspheres were measured and plotted for each line. Lines show the median with interquartile range. **C.** The clustering behavior in *KP200c* organoids is proportional to the ratio of fibroblasts to cancer cells. Bright field images of *KP200c* cancer cells co-cultured with fibroblasts at ratios of 1:1, 1:2, 1:5, and 1:10 are shown. Scale bar, 100μm. **D.** Expression of *miR-141* and *miR-200c* in *KP200c* cell lines overexpressing MSCV control or *MSCV-miR-200c/141* vector, as measured by real time PCR. U6 was used as housekeeping control. **E.** *KP* and *KP200c* cancer cells show no difference in growth rate in 2D culture. Same number of *KP* and *KP200c* cells were seeded in plates and subject to MTT assay at 24hr, 48hr and 72hr. Each condition was repeated in triplicate. Error bar, standard error. **F.** Re-expression of *miR-200c/141* reverts the invasive phenotype in *KP200c* organoids. Representative bright field images overlaid with red channel fluorescence images of *KP200c* overexpressing MSCV control or *MSCV-miR-200c/141* vector, cultured with or without tdTomato+ fibroblasts in organoid culture. Scale bar, 100μm. **G.** An Increase in CAF proliferation is observed in metastatic *KP200c* tumors. Low magnification image of low-grade, high-grade tumor, lymph node metastasis, and adrenal gland metastasis from *KP200c* animal stained for Ki67 and α-SMA. Arrows indicate proliferating CAFs that are positive for both Ki-67 and α-SMA. Scale bar, 100μm. **H.** HGF and TFGβ2 treatment induces invasive phenotype in *KP200c* tumor organoids. Phase contrast image of *KP200c* cancer cells in organoid culture untreated or treated with HGF (50 ng/ml), TGFβ2 (10 ng/ml), and a combination of HGF and TGFβ2 are shown. Scale bar, 50μm. **I.** *KP200c* cancer cell lines are able to develop lung tumors in an orthotropic allograft lung tumor model. Two *KP* and three *KP200c* cancer cell lines were transplanted into the lungs of *nu/nu* mice (1×10^6^ cells/mouse). Lungs were collected 7 weeks after transplantation and H&E images of the lungs were shown. **J. K.** *KP200c* cancer cells primed with lung fibroblast yield highly metastatic lung tumors in an orthotropic allograft tumor model. **J.** Kaplan–Meier curve of mice transplanted with *KP200c* cancer cells co-cultured with or without lung fibroblasts. Animal number N is indicated in the figure, Log rank test ** *P* =0.0062. **K.** Photos of mice 4 weeks after transplantation with *KP200c* cancer cells co-cultured with or without fibroblasts. Red arrows indicate subcutaneous metastases in mice transplanted with *KP200c* cancer cells + fibroblasts. **L.** The presence of α-SMA+ CAFs with cancer cells inside blood/lymphatic vessel of *KP200c* lung. *KP200c* primary tumor was stained for α-SMA, with the boxed area showing the presence of cancer cells accompanied by α-SMA+ CAFs inside of the blood vessel. Red arrows indicate cancer cells, green arrows indicate α-SMA+ CAFs. C: Cancer cells V: blood/lymphatic vessel. Scale bar, 100μm. **M.** α-SMA+ CAFs adjacent to circulating cancer cells in pleural lavage in a human lung adenocarcinoma patient. Blood sample collected from a lung cancer patient was stained with anti-CK19 for cancer cells and anti-α-SMA for CAFs. H&E staining and fluorescence image are shown. Scale bar, 100μm. **N.** The presence of α-SMA+ CAFs during extravasation in a human minimally invasive adenocarcinoma sample (MIA). Red arrows indicate the minimum metastatic outgrowth encased in α-SMA+ CAFs extravasating from the lymphatic vessel in the lymph node. Scale bar, 100μm V: Lymphatic vessel, C: Cancer cells, L: lymphocyte

**Supplementary Figure S4.**
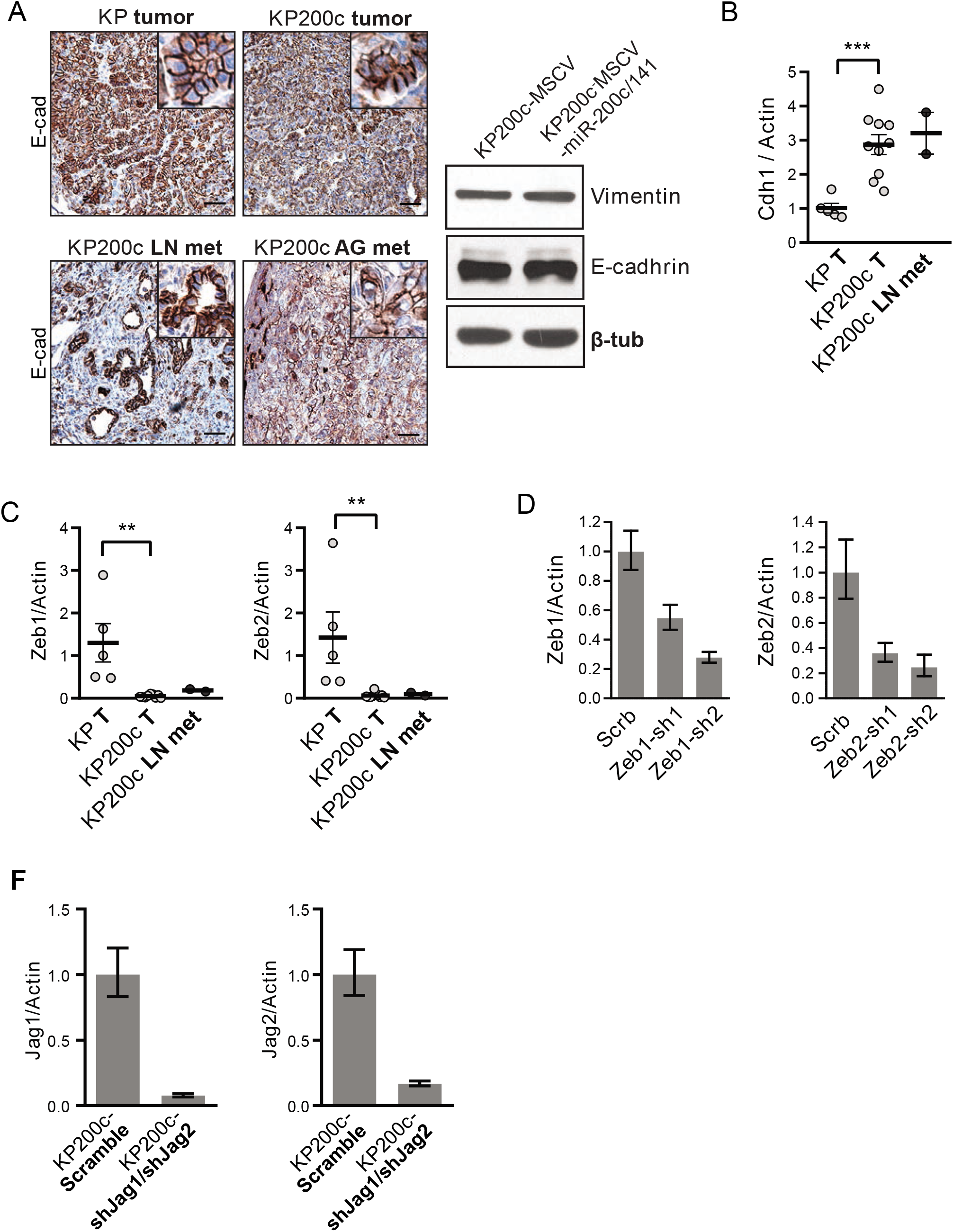
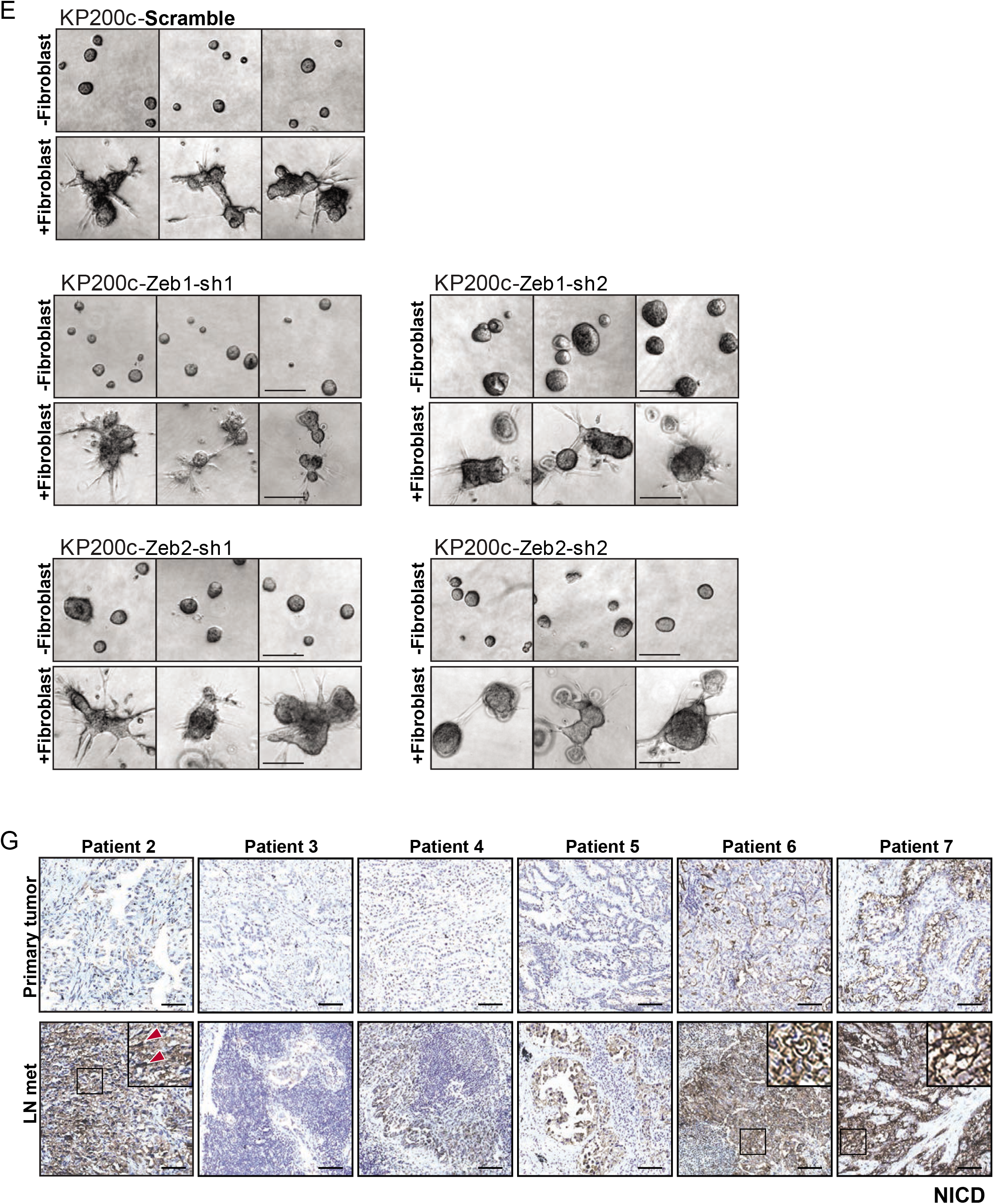
*miR-200-EMT* is not required for cancer-CAF interaction and invasive behavior. **A.** *KP200c* tumors and metastases retain epithelial characteristics. Left, *KP200c* tumors and metastases retain membrane localization of E-cadherin. Representative images of E-cadherin staining of *KP* primary tumor, *KP200c* primary tumor, *KP200c* LN metastasis, and adrenal gland metastasis are shown. Zoom in area were shown. Scale bar, 100μm. Right, expression of mesenchymal marker Vimentin and epithelial marker E-cadherin is not affected by *miR-200c/141* expression in *KP200c* cancer cells, as measured by Western blot. β-tubulin was used as endogenous control. **B.** Upregulation of E-cadherin and **C.** downregulation of *Zeb1* and *Zeb2* in *KP200c* tumor and metastases. mRNA from *KPp*rimary tumors (n=5), *KP200c* primary tumor (n=10) and LN metastases (n=2) was collected and subjected to real time PCR. The expression level was normalized to *KP* primary tumors. Error bar, standard error. *KP* primary tumor vs. *KP200c* primary tumor, *Cdh1* \*\*\**P=*0.0009, t=4.307, df=13, *Zeb1* ** *P=*0.0023, t=3.857, df=12, *Zeb2 **P=0,0055*, t=3.321, df=13, unpaired two tailed Student’s t-test. **D.** Knockdown of *Zeb1* or *Zeb2* expression in *KP200c* cell line by shRNAs. For each gene, two shRNAs were used. **E.** Knockdown of *Zeb1* or *Zeb2* does not alter invasive morphology *of* KP200c cancer cells in 3D organoid co-culture. Phase contrast images of *KP200c* cancer cells expressing Zeb1 or Zeb2 shRNAs in organoid culture with or without fibroblasts are shown. Scale bar, 100μm. **F.** Knockdown of *Jag1* and *Jag2* in *KP200c* cancer cells expressing double shRNAs against *Jag1* and *Jag2* was confirmed by real time PCR analysis. **G.** Activation of Notch in the metastases of lung cancer patients. Paired primary lung tumor and LN metastases from six lung adenocarcinoma patients were stained with anti-NICD1 antibody and nuclei were counterstained with Hematoxylin. Scale bar,100μm.

**Supplementary Table S1** This table contains the miRNA expression data as ΔCT (CT[miRNA of interest]-CT[housekeeping sRNAs]) for seven *KPT* primary tumors and four *KPT* distant metastases. Total miRNA expression level as ΔCt(*KPT*. primary tumor)+ ΔCt(*KPT*.met), fold change between *KPT* primary tumors and distant metastases as ΔCt(*KPT*.met)-ΔCt(*KPT*.primary tumor), and Student’s test *P*-value are shown.

**Supplementary Table S2** This table contains the metastasis ratio, metastasis sites and median survival time in the cohorts of *KP, KP200c* and *KP200cCKO* mice infected with high-dose Cre virus (Adeno-Cre 5×10^6^ PFU/mouse) and low-dose Cre virus (Lenti-Cre 1×10^5^ PFU/mouse).

**Supplementary Table S3** Primer sequence of real time PCR on mouse samples.

**Supplementary Table S4** Guide strand sequences for all shRNAs used in this study.

